# Mice, marmosets and humans use ambiguous local motion to stabilize gaze

**DOI:** 10.64898/2026.08.31.747047

**Authors:** Federica B. Rosselli, Marc Büttner, Annalisa Bucci, Jude F. Mitchell, Felix Franke

## Abstract

Animals need to infer properties of the environment that are only ambiguously encoded by neurons in the sensory periphery. Can these ambiguous signals drive behavior directly, or must the ambiguity first be resolved?

We traced motion signals from the retina through the nucleus of the optic tract (NOT) to reflexive eye movements, combining retinal and in vivo electrophysiology, computational modelling, and eye tracking in mice, marmosets and humans. Using stimuli that dissociate local edge motion from global, object motion, we found that responses of NOT neurons were strongly biased towards local motion. This bias propagated to behavior: eye movements followed local motion; in mice, to the point that changing the edge orientation of vertically moving objects produced horizontal eye movements. Marmosets and humans exhibited similar biases, amplified in humans when central vision was masked. Thus, the brain can transform ambiguous sensory signals directly into behavior, without first resolving the ambiguity.

## Introduction

A central problem in neuroscience is understanding how neural representations are transformed into behavior. In vision, neurons at the sensory periphery (i.e., the retina) sample only restricted parts of the environment through local receptive fields. Therefore, some properties of the environment cannot be unambiguously determined from any single local signal. As the nervous system transforms sensory signals to guide behavior, is this ambiguity resolved, or can it reach behavior?

Visual motion provides a concrete example. When an object moves through the visual scene, a small receptive field will not see the entire object but only parts of it. Within a single receptive field, only the component of motion perpendicular to the local edge can be inferred. Therefore, multiple directions of object motion are compatible with the same local measurement, a phenomenon known as the aperture problem (*1, 2*). At later stages of visual processing, this ambiguity can be resolved by combining local motion signals into a so-called ‘global’ motion representation where true object motion is accurately represented; a computationally demanding operation generally associated, in primates, with cortical visual circuits (*3–11*). The ability of cortical circuits to ‘solve’ the aperture problem does not establish that every behavior guided by visual motion depends on such a resolved representation.

Reflexive gaze stabilization provides a direct opportunity to trace the transformation of local motion signals from sensory input to behavioral response (*12, 13*). During self-motion, or when the visual environment moves relative to an observer, motion sweeps across the retina. This retinal motion elicits gaze stabilizing eye movements, known as the optokinetic reflex (OKR), that reduce retinal slip and stabilize vision (*14*).

In many mammals, the OKR for horizontal motion is mediated largely by a compact subcortical pathway centered on the nucleus of the optic tract (NOT) (*15–20*). In mice, NOT receives direct input from direction-selective (DS) retinal ganglion cells (*21–24*) which encode motion within spatially restricted receptive fields (*25–30*) (Fig. 1A, green circles). These cells can, therefore, only signal the motion component perpendicular to the local edge orientation (Fig. 1A, green arrows). Thus, this pathway begins with local, ambiguous motion signals and provides a direct link to eye-movement control. The recent identification of direction-selective ganglion cells in primate retina (*28–30*), together with the contribution of cortical motion pathways to primate gaze control (*31–36*), suggests that similar local retinal signals may contribute to gaze stabilization across species but undergo different transformations before behavior. We therefore asked whether ambiguous local motion is resolved before driving reflexive eye movements (Fig. 1B), and whether the extent to which this ambiguity reaches behavior differs between mice, marmosets and humans.

**Fig. 1:**
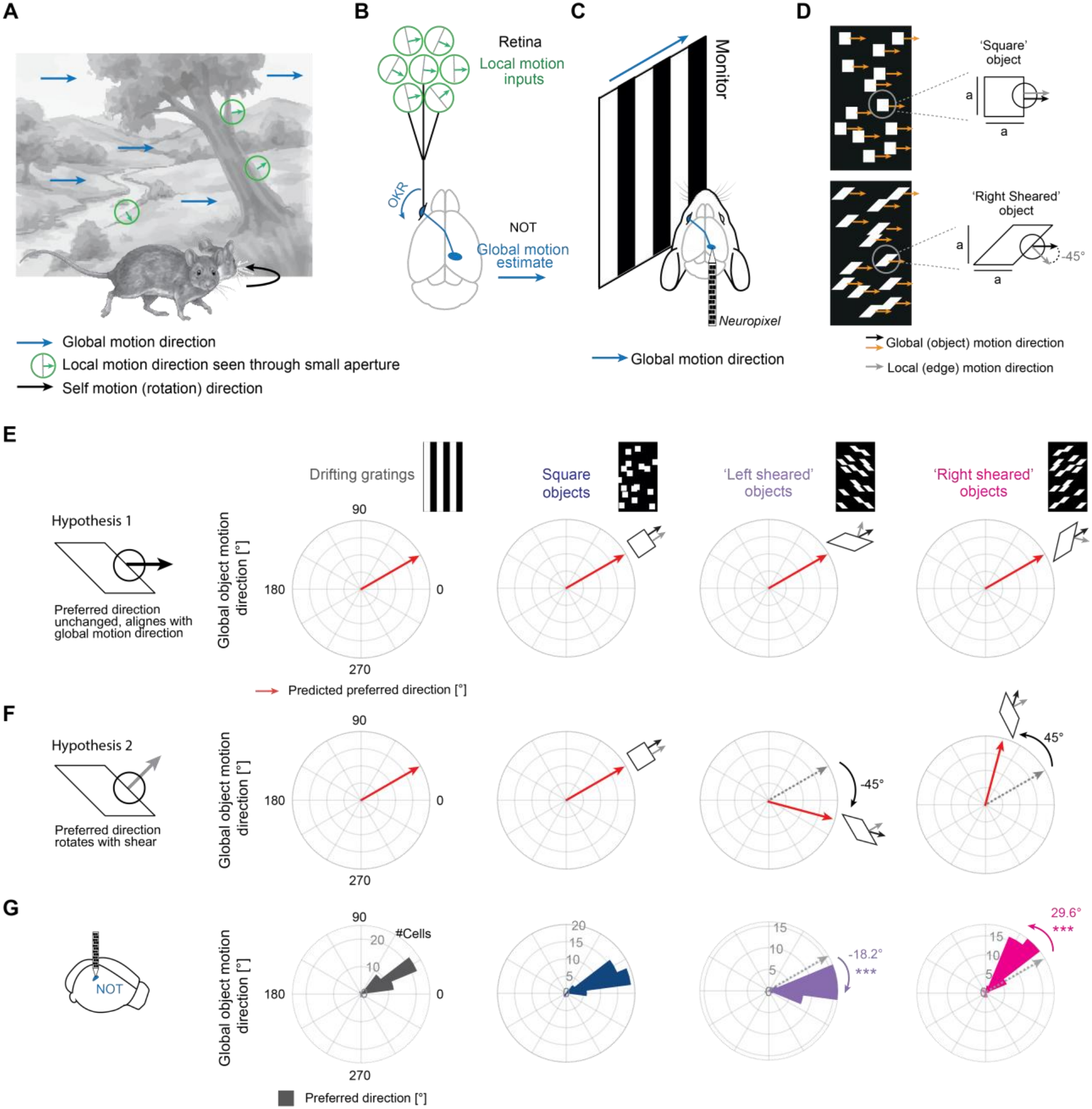
Local motion signals bias neural responses. **(A)** Global and local motion signals in a natural environment during the animal’s self-motion (head rotation). Green circles represent small apertures, through which only local motion can be estimated (green arrows). **(B)** Hypothesized computation being tested: NOT computes a global motion estimate from local motion retinal inputs to drive the OKR. **(C)** Schematics of experimental protocol for neural recordings during drifting gratings stimulation. **(D)** Left: example stimulus frames as they appear on the visual screen for square (top) and ‘right sheared’ (bottom) objects. Right: schematics of ‘square’ and ‘right sheared’ objects and their local (gray arrow) and global (black arrow) motion directions when seen through an aperture (circles). *a* is the magnitude of the square side, invariant under shearing. **(E)** Schematic depiction of predicted neural responses under Hypothesis 1 (NOT encodes global object motion). Red arrows: predicted median preferred direction as a function of motion direction of all stimulus conditions. **(F)** Schematic depiction of predicted neural responses under Hypothesis 2 (NOT encodes local edge motion). Red arrows as in **E**. Gray dashed arrows represent the native preferred direction of NOT cells, rotating when it aligns with the objects’ local motion direction. Icons represent the global object direction effectively driving the neural response. **(G)** Preferred directions distributions for n=46 cells across stimulus conditions (medians 29.2°, 20.6°, 3.4° and 50.8°). Direction (curved colored arrows), magnitude and statistical significance of the shift with respect to median preferred directions in response to square objects are reported (permutation test; left shear: P=6 × 10^−4^; right shear: P=6 × 10^−4^).

## Results

### Preferred directions of NOT cells rotate with local edge motion

To characterize how visual motion is represented in the horizontal optokinetic pathway, we recorded extracellular activity from NOT in the right hemisphere using Neuropixels probes (*37*) during passive, monocular visual stimulation of the left eye in anesthetized mice (Fig. 1C). Probe placement, and correct targeting of NOT, was verified by post hoc histology and, in Hoxd10 mice (*21*), by genetic labeling of retinal inputs to the accessory optic system (fig. S1).

We first used drifting gratings (size: 20°, speed: 45°/second; 8 directions of motion) to identify direction-selective (DS) neurons in NOT and to characterize their preferred motion directions. Only cells that met our criterion for direction selectivity (see Methods) were included in the subsequent data analyses. A substantial fraction of recorded cells (47%) exhibited robust direction selectivity (fig. S2, A, left), with preferred directions biased toward temporo-nasal motion and a slight upward component (fig. S2, A, right), consistent with previous reports (*38, 39*).

To investigate whether the mouse NOT computes a global motion estimate, we designed stimuli in which local edge orientation could be manipulated independently of object motion direction. The stimuli were composed of sheared squares, i.e., parallelograms with 45° tilted leading and trailing edges with respect to the objects’ movement direction (Fig. 1D; size: 20°, speed: 45°/second). In these ‘sheared’ object stimuli, all objects moved in the same global direction, and all were tilted either positively (‘left sheared’) or negatively (‘right sheared’). As a result, local motion signals generated along the object edges were rotated by ±45° relative to the global direction of motion.

We first compared DS responses of individual NOT cells to coherently moving square objects. Square objects evoked robust DS responses, with tuning curves closely resembling those obtained with drifting gratings (fig. S2, B, top). When the same objects were sheared to rotate local edge orientation while preserving global motion direction, tuning curves systematically rotated: clockwise for ‘left sheared’ objects, and counter-clockwise for ‘right sheared’ objects (fig. S2, B, bottom). At the population level, no loss of direction selectivity was observed across the stimulus conditions (fig. S2, C), indicating that the changes reflected a shift in tuning rather than a degradation of direction selectivity.

To interpret these effects, we compared the population tuning functions against the predictions of two alternative hypotheses. If NOT neurons encoded global object motion, preferred directions should remain unchanged for square and sheared objects regardless of edge orientation (Hypothesis 1; Fig. 1E), because the most effective stimulus would be the one in which global motion aligns with the cell’s preferred direction (black arrows in inset). In contrast, if neurons encoded local edge motion, preferred directions should rotate with edge orientation by up to −45° and +45° for ‘left’ and ‘right’ sheared objects, respectively (Hypothesis 2; Fig. 1F), because the most effective stimulus would be the one in which local edge motion aligns with the cell’s preferred direction (gray arrows in inset).

Consistent with Hypothesis 2, population responses in NOT showed systematic rotations, relative to their preferred direction, as predicted by a bias towards local edge motion (median rotations of −18.2° for ‘left sheared’ and +29.6° for ‘right sheared’; Fig. 1G). These results show that dissociating local and global motion cues produces systematic shifts in NOT cell tuning, consistent with the encoding of local (edge), rather than global (object) motion direction.

### Local edge motion biases optokinetic responses in mice

The fact that NOT does not explicitly estimate global motion does not, by itself, imply that downstream circuits fail to do so. To address this possibility, we analyzed horizontal optokinetic behavior from awake, head-fixed mice across stimulus conditions and in response to eight motion directions (Fig. 2A). If global motion was estimated beyond NOT to support the horizontal OKR, this behavior should be determined by the global direction of object motion and therefore remain unaffected by object shear. If, on the other hand, the visual system did not infer global motion to support the horizontal OKR, we should observe a specific shear-induced bias in behavioral responses, similar to that observed for neural tuning.

**Fig. 2:**
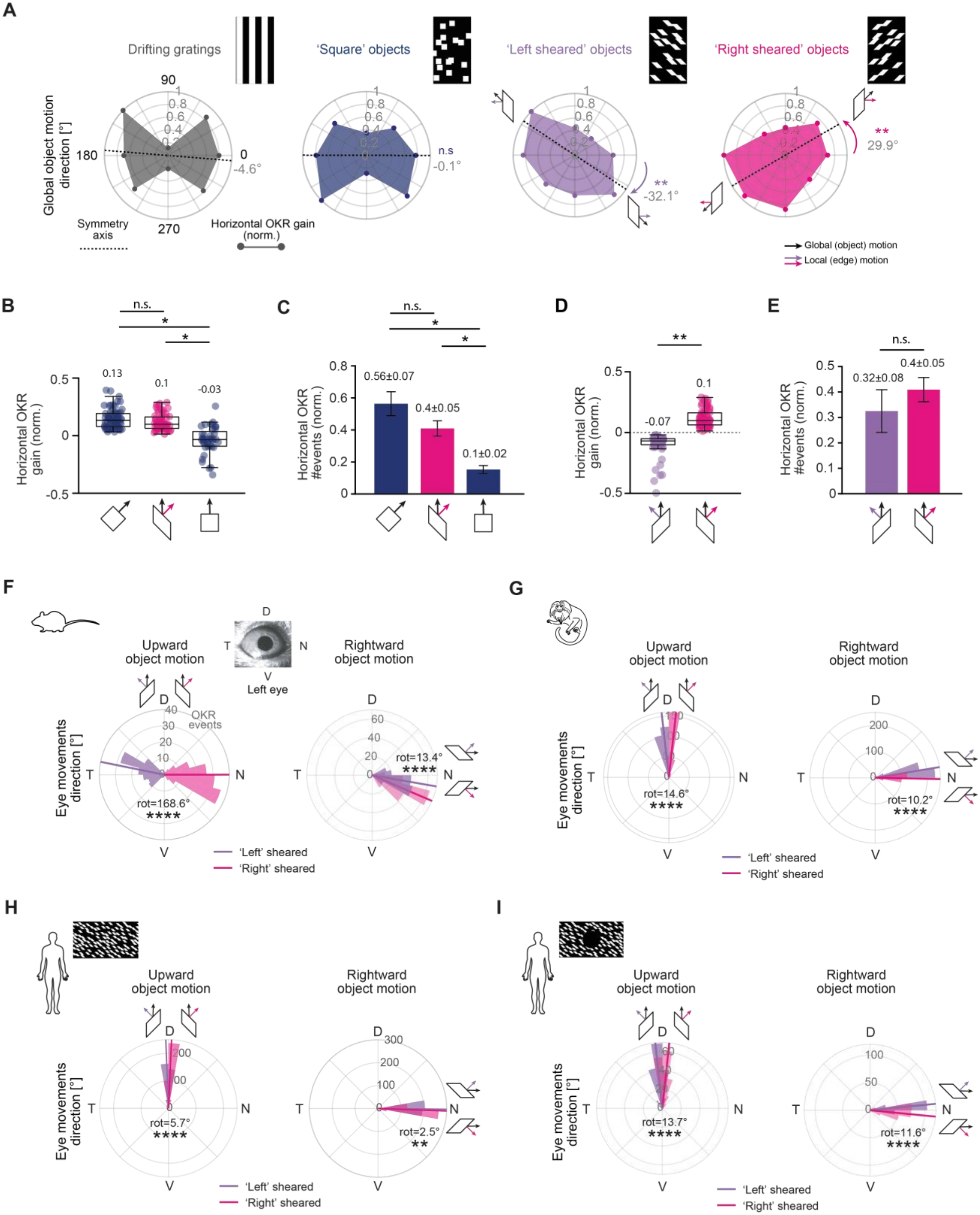
Local motion drives optokinetic eye movements away from the direction of global motion. **(A)** Distribution of normalized horizontal OKR gain values across stimulus conditions and motion directions. Dashed lines indicate the dominant symmetry axis of the gain distributions (n=6 mice). Numbers show mean angular deviation values (in degrees) of the symmetry axis from the horizontal reference axis. Statistical significance of difference in symmetry axis orientation between drifting gratings and ‘square’, ‘left’ or ‘right sheared’ objects is shown (bootstrap test on symmetry axis; P=0.65, P=0.002 and P=0.008 for ‘square’, ‘left’ and ‘right sheared’ objects, respectively). **(B)** Horizontal OKR gain values in response to square and ‘right sheared’ objects moving upward and up-right, normalized to the maximum value across stimuli. Median values are shown above the boxplots. Data points: mean gain per stimulus trial. Statistical significance of differences is shown (Wilcoxon signed-rank test, left to right: P=0.078, P=0.012 and P=0.012). **(C)** Number of horizontal OKR events per trial, normalized to the maximum mean number of OKR events per trial across stimuli. Error bars: SEM across mice (n=8). Numbers above bars: mean±SEM. Statistical difference is shown (Wilcoxon signed-rank test, left to right: P=0.109, P=0.012 and P=0.012). **(D)** Normalized horizontal OKR gain values in response to sheared objects moving upwards. Median values are shown above the boxplots. Statistical significance of differences is shown (Wilcoxon signed-rank test, P=0.008). (**E**) Normalized number of horizontal OKR events in response to sheared objects moving upwards. Numbers above bars: mean±SEM. Error bars: SEM over number of mice (n=8). Statistical significance of differences is shown (Wilcoxon signed-rank test, P=0.18). **(F)** Eye movement direction during OKR events in response to ‘left’ and ‘right sheared’ objects moving upwards (left panel) and rightwards (right panel) for n=8 mice. Histograms show data pooled across subjects. *rot* denotes the across-subjects mean angular separation between the subject-specific circular means (solid lines) for the two stimulus conditions. Statistics: linear mixed-effects model; upward motion direction: P=1.19 × 10^−132^; rightward motion direction: P=5.61 × 10^−9^. **(G)** same as **F**, but for n=2 marmoset monkeys. Statistics: linear mixed-effects model; upward motion direction: P=1.21 × 10^−23^; rightward motion direction: P=3.17 × 10^−21^. (**H**) Same as **F** and **G**, but for humans tested without a central gaze mask (n=10). Statistics: linear mixed-effects model; upward motion direction: P=6.85 × 10^−9^; rightward motion direction: P=0.002. (**I**) Same as **H**, but tested with a central gaze mask. Statistics: linear mixed-effects model; upward motion direction: P=1.76 × 10^−12^; rightward motion direction: P=4.40 × 10^−21^.

OKR responses consist of slow eye movements in the direction of stimulus motion, followed by fast resets in the opposite direction. We identified horizontal OKR ‘events’ by detecting periods of smooth eye motion between saccades (fig. S3, A) in response to stimuli presented binocularly. For each trial, we quantified the number of such events and their gain (the ratio of eye movement speed to stimulus speed) (fig. S3, B-C). Plotting horizontal OKR gain as a function of global motion direction revealed a characteristic anisotropic pattern for both gratings and square objects, resembling a sideways figure-eight with substantially higher gain for horizontal than vertical motion (Fig. 2A, gray and blue). Strikingly, this gain pattern rotated for sheared objects (Fig. 2A, purple and magenta) in the same direction as the tuning curves in the NOT neural population (Fig. 1G, purple and magenta). The rotation was substantial (-32.1° and +29.9° for ‘left’ and ‘right sheared’ stimuli, respectively), and shifted the gain maxima toward the stimulus directions for which the local edge motion, rather than global object motion, was most strongly aligned with the horizontal axis (Fig. 2A, purple and magenta, object conditions in inset).

The analysis so far did not reveal whether the eye movement direction was also affected by the shear. To test this, we focused on upward-moving ‘right sheared’ objects, a condition in which global motion was upward whereas local edge motion was up-right, i.e., toward the preferred direction of NOT neurons. We asked whether horizontal OKR responses to this condition would resemble those elicited by square objects moving upward, which matched the global motion direction, or rather those elicited by square objects moving up-right, matching the local edge motion direction.

Behavioral responses matched the prediction that eye movement direction follows local, rather than global, motion. Horizontal OKR gain values evoked by upward-moving ‘right sheared’ objects closely resembled those evoked by up-right moving square objects, despite the unchanged upward object motion (Fig. 2B). The same pattern was observed for the number of horizontal OKR events (Fig. 2C): upward-moving square objects elicited very few horizontal OKR responses, whereas upward-moving sheared objects produced robust responses.

Because upward-moving sheared objects contain a local motion component to the up-left or up-right (Fig. 2D, colored arrows), they provide a stringent test of whether local motion can drive the horizontal component of the OKR. Upward-moving sheared objects elicited more horizontal than vertical OKR events as compared to the other stimuli (fig. S4A). Moreover, upwards-moving ‘left’ and ‘right sheared’ stimuli produced horizontal OKR gains of opposite sign (Fig. 2D) and substantially more horizontal OKR events than upward-moving square objects (Fig. 2E, compare with rightmost bar in Fig. 2C). This suggested that the upwards moving sheared objects elicited eye movement in different directions depending on shear direction.

To test this, we analyzed the eye movement direction in response to upwards (Fig. 2F, left) and rightwards (Fig. 2F, right) stimulus motion. In stark contrast with symmetrical stimuli (gratings and square objects), vertically moving ‘left’ and ‘right sheared’ objects produced horizontal eye movements in nearly opposite directions (fig. S5), separated by an angular shift of 168.6° (Fig. 2F, left). By comparison, the angular separation between sheared objects moving rightward was much smaller (13.4°; Fig. 2F, right), indicating that the impact of shear on eye movement direction depended strongly on the global, object motion direction.

Together, these results show that optokinetic behavior is driven primarily by local motion signals and is strongly biased toward their horizontal component, to the point that vertically moving sheared objects triggered horizontal eye movements, whose direction inverted with the direction of the shear.

### Local edge motion biases optokinetic responses in marmosets and humans, and bias strength increases when central vision is masked

To test whether the observed effect of local motion on behavior generalized across species, we analyzed OKR behavior in marmoset monkeys freely viewing the same stimuli shown to mice scaled to reflect the smaller receptive field sizes (object size: 1°). For rightward motion, marmosets exhibited a qualitative bias toward local edge motion similar to that observed in mice. For upward motion, however, the bias was substantially reduced, although it remained stronger than for rightward motion (Fig. 2G).

We next tested whether this effect generalized to humans, viewing the same stimuli shown to mice and marmosets, scaled for human viewing (object size: 5°). While humans exhibited the same qualitative bias toward local edge motion as mice and marmosets, the effect was substantially smaller, particularly for upward direction (Fig. 2H). We hypothesized that the weaker bias in primates arose from a contribution of high-acuity central vision present in foveated species, and involving cortical motion processing, attenuating an otherwise local-motion computation. This predicted that masking central visual stimulation should increase the bias to local motion and shift human responses towards the mouse phenotype. To test this possibility, we presented human participants with the same stimulus set, but this time with a circular black mask spanning 33° at the center of the visual field. To prevent participants from viewing the stimulus using their central retina, they were instructed to maintain their gaze within the mask, and we removed eye movements that extended beyond the mask from the analysis. Masking the central visual field (Fig. 2I) markedly increased the influence of local edge motion on eye movement direction (Table S6), suggesting that central visual input attenuates, though does not eliminate, the influence of local motion on reflexive eye movements in humans. Just like marmosets, and in contrast to mice, upward and rightward motion directions elicited a local-motion-driven angular separation of similar magnitude in eye movement direction (marmosets: 14.6° and 10.2°; humans with central mask: 13.7° and 11.6°, respectively; Fig. 2I). The direction of the local motion-driven bias was consistent across all three species, and across both masked and unmasked conditions in humans (fig. S6). Together, these results show that local motion signals influenced reflexive eye movements in primates, and, in humans, particularly when central visual input was masked. However, the quantitative differences between humans and mice were not fully abolished by masking.

### Anisotropic gain transforms local motion signals into optokinetic eye movements

To explain how biased local motion signals are transformed into eye movements, and explain the quantitative differences between species, we developed a simple phenomenological model. In particular, we sought to explain two key features of optokinetic behavior revealed in Fig. 2: the strong horizontal eye movements elicited by upward-moving sheared objects, and the striking difference in eye-movement rotation between upward- and rightward-moving stimuli.

The inputs to the model were the global, object stimulus motion direction, and the local motion direction, defined by the local edge orientations. The model then related these stimulus directions to eye movement directions using two parameters. A ‘bias strength’ parameter described how strongly the local motion direction influenced the eye movements as opposed to the global motion direction. A bias strength of 100% meant that the local signals fully determined the eye movement directions, whereas a bias strength of 0% would indicate that only the global motion direction was driving eye movements. A second parameter, the ‘gain ratio’, described the ratio between the influence of the vertical motion component to the horizontal motion component. We considered two alternative models: in the *isotropic gain* model, the gain ratio was set to one, and thus horizontal and vertical motion components contributed equally to eye movement direction (Fig. 3A, left). In the *anisotropic gain* model, horizontal components were weighted differently than vertical components. For this model, and consistent with our (fig. S4A) and previous results (*21, 22, 40*) (Fig. 3A, right), we modeled the gain of the horizontal component to be stronger than the gain of the vertical component.

**Fig. 3:**
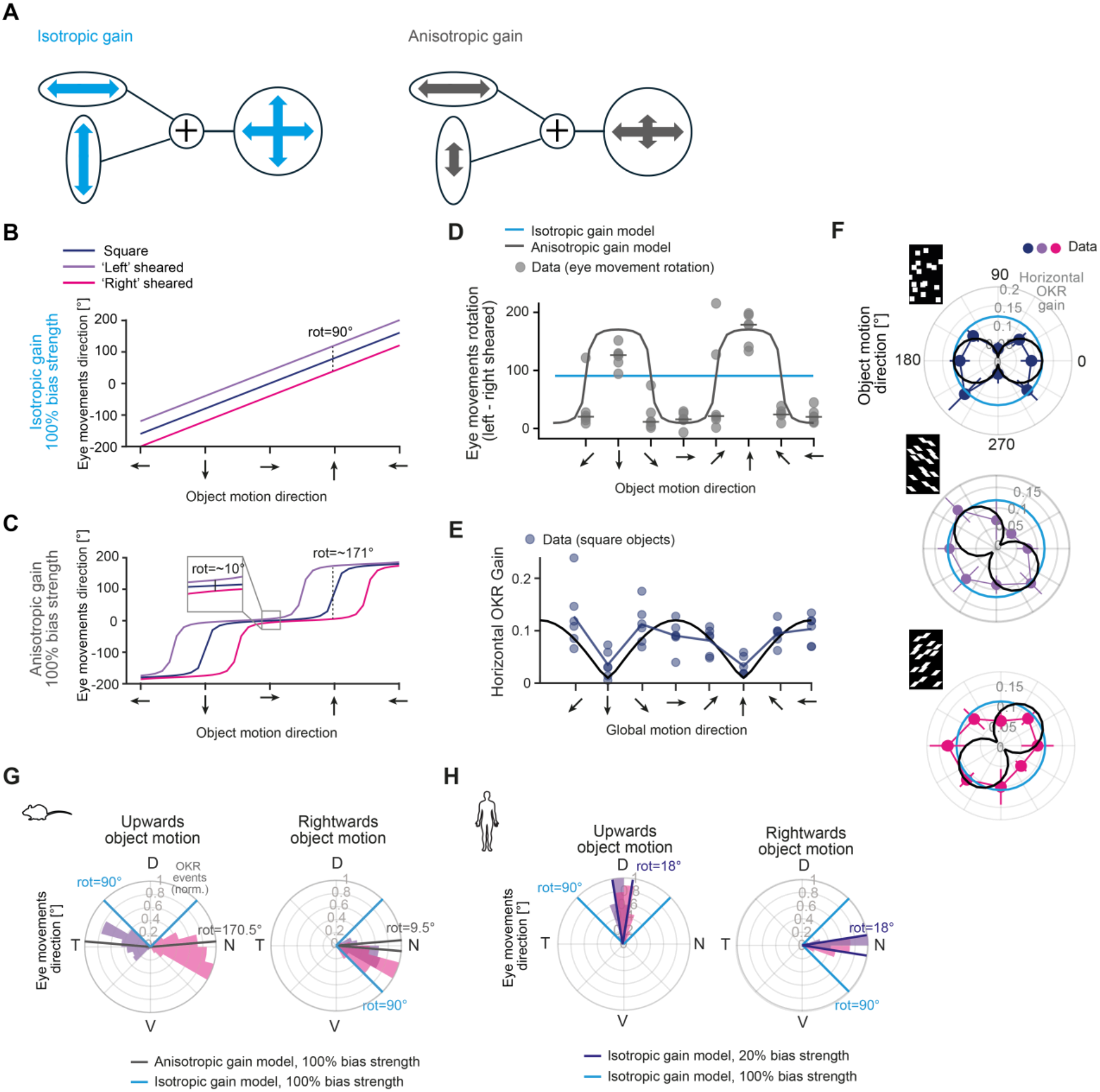
Anisotropic gain and bias strength warp local motion signals into optokinetic eye movements. **(A)** Schematics of isotropic and anisotropic gain models, where horizontal and vertical motion components are weighted equally and unequally, respectively, and are then multiplied by the strength of the bias. **(B)** Eye movement directions in response to square and sheared objects as predicted by an isotropic gain model with 100% bias strength, and angular separation (rot) between them, as a function of object (global) motion direction. **(C)** Eye movement directions as predicted by an anisotropic gain model with 100% bias strength; same structure as **B. (D)** Eye-rotation angles predicted by the two models as a function of object motion direction (continuous lines). Data points show rotation angles between responses to ‘left sheared’ and ‘right sheared’ objects at each object motion direction (n = 6 mice). **(E)** Gain distributions predicted by the two models (light blue line and dark gray lines) as a function of global motion direction. Measured OKR gain values in response to square objects (n=6 mice) are overlaid. Blue continuous line: mean of the gain values across motion directions. **(F)** Model- and data-derived OKR gain as a function of global motion direction; same data as in **E**, fig. S7, A-B and Figure 2A (normalized), shown in polar coordinates for square, ‘left’ and ‘right sheared’ objects, and where each data point is the mean±SD gain value at each global motion direction. **(G)** Model-predicted eye movement directions and rotation angles in response to ‘left’ and ‘right sheared’ objects stimuli moving upwards and rightwards for mice. Normalized data from Fig.2F is overlaid. **(H)** Same as **G** but for humans with a central gaze mask. Normalized data from Fig.2I is overlaid.

Under the isotropic gain model, eye-movement direction varied linearly with global motion direction for all stimuli (Fig. 3B). For square objects, the eye movement direction aligned with the identity line (Fig. 3B, blue line), whereas the lines for ‘left’ and ‘right sheared’ objects were shifted symmetrically above and below the identity (Fig. 3B, purple and magenta). This predicted fixed angular separations (rotations) of the eye movement direction between stimulus classes across all motion directions. This model therefore predicted similar rotations between ‘left’ and ‘right sheared’ stimuli for upward and rightward motion.

By contrast, the anisotropic gain model produced markedly different predictions (Fig. 3C). Because horizontal components were amplified relative to vertical ones, the relationship between global motion direction and eye-movement direction became highly nonlinear. For horizontal motion, predicted eye-movement directions for square and sheared objects were close, but not identical (inset in Fig. 3C), whereas for vertical motion they diverged strongly. This geometry predicted a near-180° separation between eye movements evoked by upward-moving ‘left’ and ‘right sheared’ objects, but only a small separation for rightward-moving stimuli.

We first tested these predictions against mouse behavioral data. Across motion directions, the anisotropic gain model accurately reproduced both eye-movement rotation (Fig. 3D) and OKR gain (Fig. 3E; fig. S7, A-B), whereas the isotropic model failed to capture the observed responses (Fig. 3D, light blue line).

Polar representations of gain for square and sheared stimuli further showed that the anisotropic model captured both the shape and orientation of the observed gain distributions: overlaying model predictions onto the behavioral polar plots from Fig. 2A confirmed that the anisotropic gain model accurately reproduced the full pattern of OKR gains across stimulus motion directions observed in mice (Fig. 3F).

We next quantified the angular difference between eye movements evoked by ‘left’ and ‘right sheared’ objects as a function of global motion direction. The isotropic gain model predicted a constant angular separation for both upwards and rightwards object motion (Fig. 3G, left and right, blue lines), whereas the anisotropic model predicted a strong dependency of angular separation on motion direction (Fig. 3G left and right, dark gray lines). Behavioral data was consistent with the anisotropic prediction, with large separations for upward motion and minimal separations for rightward motion (Fig. 3G, histograms).

The model illustrates how local-motion bias and gain anisotropy jointly shape optokinetic behavior. Gain anisotropy determined how local motion was transformed into eye movements, whereas bias strength determined the extent to which local edge motion influenced eye-movement direction. Whereas mouse behavior was accounted for by a strongly anisotropic gain model with 100% bias strength, human behavior was better explained by an isotropic gain model with substantially weaker local-motion bias (20%; fig. S7, C), consistent with the smaller angular separations observed experimentally (Fig. 3H).

Together, these findings show that a simple gain model combining local-motion bias with anisotropic weighting accurately accounts for optokinetic behavior across stimulus conditions and species.

### NOT cells encode motion through spatially localized, elongated receptive fields clustered along the horizontal meridian

Having characterized how local-motion bias and gain anisotropy shape eye movements, we returned to the mouse retina-NOT pathway to ask where in visual space these local signals are sampled from and how they are combined.

To determine how NOT neurons sample visual space, we estimated their receptive fields using a random motion-based mapping paradigm (*41*), consisting of a random moving objects (RMO) stimulus (Fig. 4A, left). This method quantified how often local object motion at a given display location preceded neural spiking. This approach yielded robust receptive field estimates for the majority (∼86%) of DS NOT cells, in stark contrast with pixel-contrast-based reverse correlation with white noise (*42*), which yielded reliable receptive field estimates for only half (∼48%) of DS neurons (fig. S8, A-E).

**Fig. 4:**
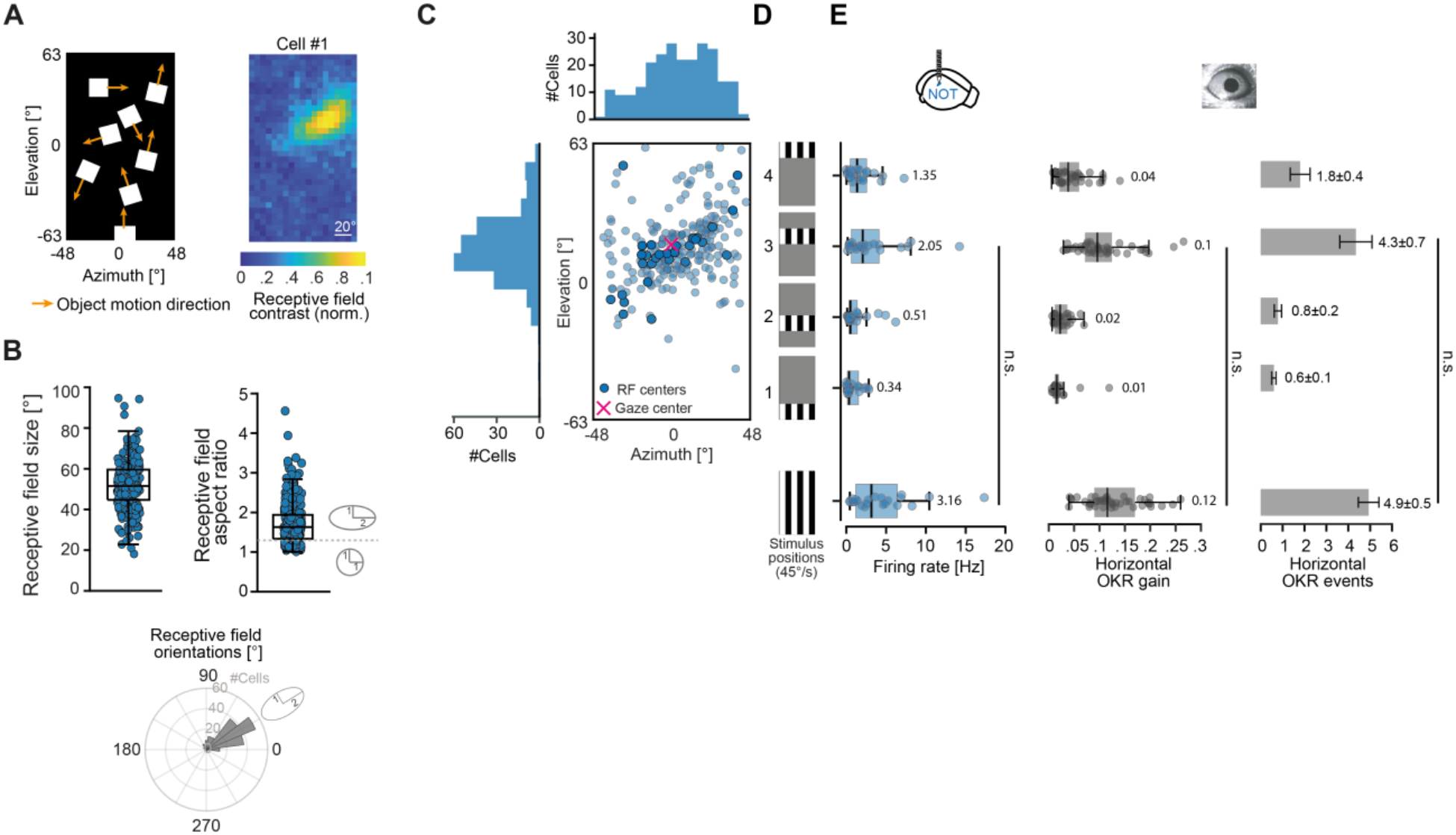
Reflexive gaze stabilization is driven by localized, spatially clustered receptive fields. **(A)** Left: depiction of a stimulus frame for the random moving objects (RMO) stimulus on visual screen coordinates; orange arrows indicate motion direction of each object. Right: one DS cell receptive field, plotted as a function of visual screen coordinates: color codes the normalized receptive field contrast. **(B)** Receptive field sizes (left, median 51.5°), aspect ratio (right, median 1.6; dashed line marks elongation threshold, schematics show examples of Gaussian fits for a round receptive field, where both axes have length 1, and an elongated receptive field, where one axis has double length) and orientation distributions (bottom, median 35.3°) for n=243 DS cells. **(C)** Receptive field positions plotted as a function of monitor coordinates the same cells in B (blue dots), and as distributions in the azimuth and elevation axes (histograms). Darker blue dots represent the cell population depicted in Fig. 1G. **(D)** Schematics of stimulus location when rightwards drifting gratings were shown as 30° stripes at 4 positions in random order (1-4), and when shown as full-field conditions. **(E)** Left: responses of n=22 DS cells as a function of stimulus position. Median firing rates are shown. No statistical difference between median firing rate at full field and stimulus position 3 (permutation test, P=1.000). Middle: horizontal OKR gain values as a function of stimulus position. Data points are mean gain value per trial across mice (n=6). Numbers above boxplots: median gain values per stimulus position. No statistical difference between full-field and position 3 (Wilcoxon signed-rank test, P=1.000). Right: horizontal OKR events (mean/trial) across mice (n=6) and stimulus positions. Error bars: SEM across mice (n=6). No statistical difference between full-field and position 3 (Wilcoxon signed-rank test, P=0.468).

Receptive fields estimated with this method were spatially localized and markedly elongated (example cell in Fig. 4A, right). Receptive field sizes spanned a wide range (∼18° to ∼94°), with a median size of 51.5° (Fig. 4B, top left). Most receptive fields exhibited an elongated aspect ratio (Fig. 4B, top right), and were systematically tilted slightly upward from the horizontal axis, and aligned with the cells’ median preferred motion directions (Fig. 4B, bottom). Similar receptive field shapes were observed in non-direction selective (non-DS) NOT cells (fig. S8, F-H), indicating that elongated spatial sampling is a general organizational feature of NOT. This geometry was not attributable to head orientation, as eye position, including ocular torsion, is actively stabilized by the vestibulo-ocular reflex (*43, 44*). We confirmed that this coupling between head and eye movements is active even under anesthesia (fig. S9, A-B).

At the population level, NOT receptive fields were not uniformly distributed across visual space. Instead, they clustered along the horizontal meridian, near the center of the visual field (Fig. 4C). When receptive field positions from multiple recording sessions, i.e. from electrode insertions at different locations, were aligned in anatomical coordinates (anteroposterior, AP; mediolateral, ML), this bias was observed consistently across recording sites, with receptive field centers from different penetrations occupying overlapping regions of visual space (fig. S9, C-F). Together, these observations indicate that the clustering along the horizontal meridian is not explained by uneven anatomical sampling, but reflects a property of NOT population organization. To test whether this spatial sampling constrains functional responses, we next examined neural and behavioral sensitivity to motion presented at different visual locations. We compared responses to full-field drifting gratings with responses to gratings restricted to non-overlapping horizontal stripes of equal visual area (Fig. 4D). NOT firing rates were maximal when gratings overlapped the region of visual space corresponding to the population receptive field cluster and were strongly reduced for gratings presented elsewhere (Fig. 4E, left). Optokinetic behavior showed the same dependence as neural activity on the visual stimulus spatial location: eye movements were not only elicited by full-field, global motion but equally well by motion localized within the receptive field-dense region (Fig. 4E, center and right). These results show that full-field motion is not required to engage NOT neural activity, nor behavioral responses; rather, localized horizontal motion near the horizontal meridian is sufficient.

### Linear pooling with local gain control accounts for NOT responses

NOT neurons sampled large, elongated regions of visual space, yet their directional responses remained biased toward local edge motion. We therefore asked whether linear pooling of spatially distributed local retinal motion signals, which by itself cannot recover global object motion (*1, 45, 46*), could account for both properties.

We constructed a two-stage model comprising a fixed retinal stage that generated local motion signals and a NOT stage whose integration of those signals was fitted to neuronal responses (Fig. 5A). To inform the retinal stage, we recorded retinal ganglion cells from isolated mouse retina using high-density multielectrode arrays (HDMEAs (*47*)) (fig. S10). We estimated retinal receptive fields with the same motion-based mapping approach used for NOT, with objects scaled down to 7° to resolve the smaller retinal receptive fields. Direction tuning functions were estimated using a moving bar stimulus (fig. S10C). Direction-selective retinal ganglion cells (DS-RGCs) exhibited broad direction tuning (fig. S10D) consistent with previous reports (*21, 22*). Receptive fields of DS-RGCs, including DS-RGCs preferring temporal motion, were about five times smaller than NOT receptive fields (median ∼11° vs. 51.5°; fig. S10, B and E, left). They were also approximately circular (median aspect ratio 1.1; fig. S10, E, right).

**Fig. 5:**
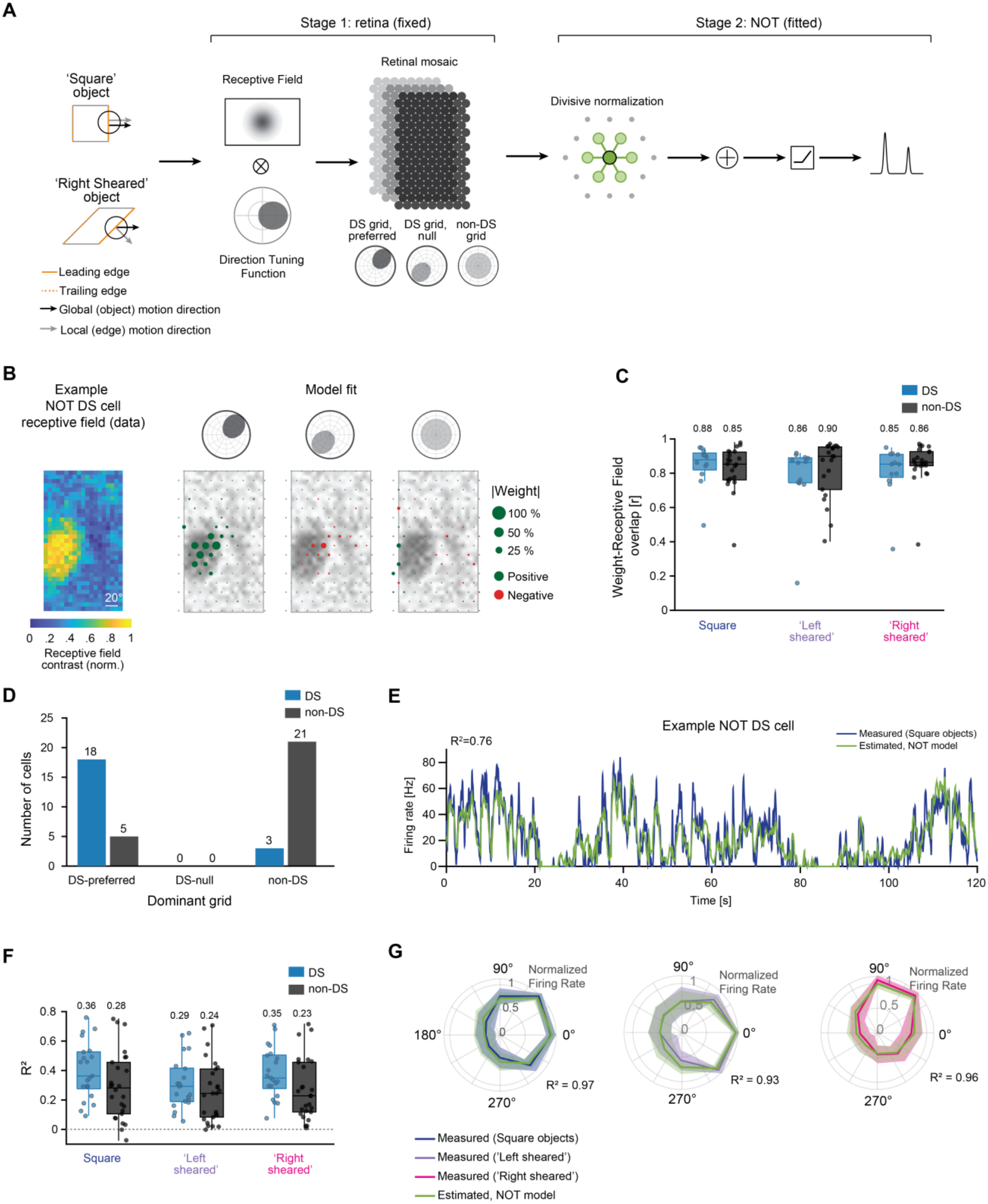
Linear pooling with local gain control accounts for NOT responses. **(A)** Schematics of the NOT model. Left: leading and trailing edges of a ‘square’ and a ‘right sheared’ object. Within a receptive field (circle), the edge of a sheared object moves perpendicular to its orientation (local edge motion direction), at 45° to the object’s motion (global motion direction). At the retina stage, local motion detectors combine a Gaussian receptive field with a von Mises direction tuning function and are arranged in three hexagonal grids representing three retinal input classes: DS cells tuned to the NOT preferred direction (DS grid, preferred), DS cells tuned to the opposite direction (DS grid, null) and non-DS cells (non-DS grid), each grid once for the leading and once for the trailing edge. Divisive normalization: each detector’s output is divided by the non-DS input pooled over its own grid node and the neighboring nodes (green). The model NOT cell computes a weighted sum of the inputs and passes the sum through a rectifying nonlinearity to predict the firing rate. **(B)** Receptive field of an example DS NOT cell (color codes the normalized receptive field contrast), and the cell’s model weights on the DS-preferred, DS-null and non-DS grids for square objects; the leading- and trailing-edge weights of each grid node are summed. Dot area is proportional to the weight magnitude, scaled to the largest node weight of the three maps; green, positive; red, negative. Gray shading: the cell’s receptive field; gray points: grid nodes. **(C)** Pearson correlation distribution quantifying the overlap between NOT cells’ receptive fields and detector weight locations. Only cells with a receptive field quality index Qi > 0.3 are included (DS, blue, n = 12; non-DS, black, n = 22). Median values are annotated as numbers above boxes. Boxes: median and quartiles; whiskers: most extreme values within 1.5 × IQR; dots: individual cells. **(D)** Number of cells per dominant grid, i.e. the grid carrying the largest summed absolute weight, for DS (blue, n = 21) and non-DS (black, n = 26) cells in response to square objects. **(E)** Measured firing rate (blue) and cross-validated model prediction (green) of the example cell in B during a continuous, 10-minute presentation of square objects (120 s shown). R^2^ = 0.76: cross-validated coefficient of determination over the whole trace. **(F)** Cross-validated R^2^ per cell for DS (blue, n = 21) and non-DS (black, n = 26) NOT cells in response to square, ‘left sheared’ and ‘right sheared’ objects. Median values are annotated as numbers above boxes. Boxes as in C; dotted line: R^2^ = 0. **(G)** Population mean direction tuning of measured and predicted (green) responses of the DS cells (n = 21) to square (dark blue), ‘left sheared’ (purple) and ‘right sheared’ (magenta) objects, the model was fitted separately to each condition. Each cell’s measured and predicted tuning curves were normalized to the peak of its measured curve; Shaded regions: SD of the measured and of the model responses across cells. R^2^: coefficient of determination of the model for the measured population-mean tuning curve.

We modeled DS-RGCs as local motion detectors that respond to the leading or trailing edges of moving objects, reflecting known properties of DS-RGCs (*14*). Each detector combined a circular Gaussian receptive field with a von Mises direction tuning function, parameterized to reproduce the key features of the measured DS-RGCs: small receptive fields relative to NOT and broad direction tuning (Fig. S10). Importantly, the parameters of this ‘retinal stage’ were held fixed throughout NOT model fitting.

Modeled retinal ganglion cells formed three classes of input to the second, NOT, stage: DS-RGCs tuned near the preferred direction of the NOT population, DS-RGCs tuned to the opposite direction, and non-direction-selective (non-DS) cells. Each class was represented by separate retinal mosaics responding to leading and trailing object edges, giving six mosaics spanning the visual field (Fig. 5A). We first tested a NOT stage that combined RGC outputs through weighted linear summation. Input weights and an intercept were fitted to each neuron’s firing rate using sparse regularization, without supplying the measured NOT receptive-field location or shape. A fixed output rectification prevented negative firing-rate predictions. Models were fitted separately for square, left-sheared and right-sheared objects, and evaluated on held-out temporal blocks within each condition. This linear pooling model predicted the smoothed firing-rate responses of DS NOT neurons with median cross-validated R^2^ values of 0.29, 0.21 and 0.30, respectively (fig. S11B, ‘Lasso’).

We next tested whether nonlinear integration improved predictive performance while retaining the same fixed retinal stage. In one variant, each RGC output was divided by a local signal pooled from neighboring non-DS RGCs before weighted summation. This divisive normalization introduced no additional parameters fitted to NOT responses, and carried no direction information (*48*); NOT-stage weights and intercept were estimated using the same regularized fitting procedure. In a second variant, boosted regression trees replaced the weighted summation stage. Divisive normalization increased median cross-validated R^2^ to 0.36, 0.29 and 0.35 across the three stimulus conditions, whereas boosted trees yielded corresponding values of 0.37, 0.26 and 0.34 (fig. S11B). Thus, local normalization improved prediction, while the more flexible tree model provided no consistent additional advantage. Comparisons for non-DS NOT neurons are shown in fig. S11C. We therefore used the ‘divisive normalization’ model to examine how retinal inputs accounted for NOT response properties. The model fit assigned high input weights at visual locations that closely matched the experimentally estimated receptive fields, although those receptive fields were not used to constrain model fitting (Fig. 5B–C). Spatial correspondence was also observed for weights of the linear model and from tree feature importance (fig. S11D–E). In the normalized model, DS NOT neurons predominantly assigned weight to the RGC class with compatible direction preference, whereas non-DS NOT neurons predominantly assigned weight to non-DS RGCs (Fig. 5D). The divisive normalization model captured temporal fluctuations in NOT firing rates (Fig. 5E–F) and closely reproduced the population direction tuning for each stimulus condition, including the shear-dependent shifts (Fig. 5G; population tuning R^2^ = 0.93–0.97). Together, these results show that spatial pooling of local retinal motion signals accounts for key features of NOT responses. Local gain control improves this account, while the nonlinear tree model tested here provided little further predictive benefit.

## Discussion

By placing local and global motion signals in conflict and tracing them through each stage of the reflexive gaze-stabilization circuit in mice, marmosets and humans, we found that ambiguous local motion can reach behavior without first being resolved by the visual system. In mice, motion direction tuning in NOT was strongly biased toward local edge motion, and this bias was preserved in optokinetic behavior rather than corrected downstream.

Marmosets and humans showed a similar, albeit weaker, local-motion bias, and the differences between mice and humans were captured by a phenomenological model with two parameters: the anisotropy between horizontal and vertical gain, and the strength of the local-motion bias. The presence of this bias in all three species, together with the recent identification of direction-selective ganglion cells in primate and human retina (*28–30*), suggests that it is a general feature of reflexive gaze stabilization rather than a species-specific property of the mouse retina-NOT pathway. In humans, the bias was strongest when central vision was masked. In primates, which are foveated and in which cortical motion areas contribute to object-motion processing and gaze control (*4, 7, 11, 31–33*), central vision may therefore provide high-resolution information about object motion that counteracts ambiguous local signals from the periphery. Although the direction of the bias was consistent across subjects, its magnitude varied, potentially reflecting differences in attentional engagement, which strongly modulates optokinetic responses to peripheral motion (*49*).

Two further findings in mice elucidate how the bias arises. First, neither NOT activity nor the horizontal OKR required motion across the full visual field; instead, the pathway preferentially sampled a restricted region near the horizontal meridian. This suggests that the circuit is specialized for the horizontal retinal slip generated by translation or common rotational movements (*50*), for which horizontal compensatory eye movements are particularly effective (*51*). Second, a model in which NOT cells linearly pool locally normalized retinal motion signals accounted for the defining properties of NOT responses. Because the normalization signal was local and carried no direction information, it could rescale local inputs but not combine motion directions across the visual field into a global estimate. Thus, in mice, reflexive gaze stabilization, a behavior commonly framed as a global-motion problem (*12, 15, 52–54*), can operate without an explicit estimate of global object motion.

A useful way to formalize these findings is to distinguish strong from weak solutions to the aperture problem. A strong solution recovers global object motion by combining ambiguous local measurements into a single, geometrically consistent velocity estimate through nonlinear computations such as intersection-of-constraints or least-squares integration (*1, 3, 4, 45, 46*). Linear summation cannot, in general, achieve this, because the relationship between local image motion and object motion is itself nonlinear. A weak solution leaves the ambiguity unresolved, but nevertheless produces an appropriate output when the distribution and weighting of local signals happen to align with the behaviorally relevant direction. Many conventional motion stimuli fall into this category (*55, 56*): because their local cues are symmetrically distributed around the global direction, pooling local motion yields responses aligned with object motion, allowing local pooling to masquerade as global-motion inference. This may explain why reflexive motion systems have often appeared to compute global motion. Stimuli that displace local cues to one side of the global direction, such as our sheared objects, separate the two regimes, and under these conditions both NOT activity and optokinetic behavior followed the dominant local motion rather than the object’s true direction. The retina–NOT–OKR pathway therefore operates in the weak regime.

Type II plaids, in which both component velocities lie on the same side of the pattern direction, pose an analogous test for perception, and human observers can be strongly biased toward component-based solutions, particularly at short presentation durations (*57, 58*). One possibility is that rapid motion responses are initially dominated by local signals, whereas additional processing associated with central vision and cortical pathways subsequently constrains motion estimates toward object-consistent solutions.

Functionally, a weak solution may be well matched to the demands of gaze stabilization during self-motion. The primary goal of the OKR is to minimize retinal slip rapidly, not to infer the trajectories of individual objects, and under natural viewing conditions local and global optic flow will often be aligned closely enough for pooled local motion to provide an effective stabilizing command. Explicitly reconstructing object motion would add computational complexity without necessarily improving stabilization. The errors revealed by sheared objects may therefore be the predictable cost of a system optimized for speed, robustness and the statistics of its usual sensory input.

## Supplementary Material

Materials and methods

Figs. S1 to S11

Tables S1 to S8

## Materials and methods

### Animal preparation, mice experiments

The Institute of Molecular and Clinical Ophthalmology Basel (IOB) hosted the animals in a specific pathogen free facility. The housing conditions were controlled with a 12:12 light-dark cycle, room temperature in the range of 20-26° C and relative humidity in the range of 45%-65%. Water and food were provided *ad libitum*. Researchers and veterinarians daily checked the health and welfare general conditions of the animals according to the Swiss animal protection law.

All experiments were performed on adult wild-type (WT; C57BL/6; Charles River Laboratories; strain code 027; RRID:IMSR_CRL:027) and Hoxd10 (Tg(Hoxd10-EGFP)LT174Gsat/Mmucd; MMRRC stock no. 032065-UCD; RRID:MMRRC_032065-UCD (*21*)) mice, male and female, with age ranging from 2 to 8 months (n=8 for psychophysics, n=22 for neural recordings, n=14 for HDMEA recordings). On the day of the recording, a mouse was anesthetized using a subcutaneous (SC) injection of Fentanyl-Medetomidine-Midazolam (FMM; Fentanyl, Janssen, 0.05 mg/kg; Medetomidine, Virbac AG, 0.5 mg/kg; Midazolam, Sintetica, 5 mg/kg; in 0.9% saline) and placed on a heating pad to prevent anesthesia-induced hypothermia. Eye lubricant was applied throughout the procedure to prevent corneal dehydration. The scalp was shaved using hair clippers and disinfected with Iodine solution. Local anesthesia (Lidocaine (0.1-0.2%)) was administered subcutaneously prior to incision. A ∼5 mm scalp section was removed via microscissors. The periosteum was removed and the scalp edges were fixed to the skull using surgical glue (Histoacryl, *Braun*) to avoid interference with the implant. The skull was then etched using a microcurette and a drop of hydrogen peroxide to ensure cement grip. Finally, a custom-built aluminum headpost was affixed to the skull using super-glue (Loctite 403, *Conrad*) and dental cement (Super Bond C&B, *MPE Medical GmbH*). Animals undergoing psychophysics (n=8) were left to recover in their home cage for three days before the habituation period started. In this period, animals were habituated to manipulation and head-fixation in the recording setup for a minimum of 1 to a maximum of 5 days before eye-tracking experiments started.

For animals undergoing neural recordings, on the day of the recording, a ∼3×3mm craniotomy was performed with a 0.7 mm burr dental drill (AEU-25 and AHP-64, *Altmann Dental*) above the approximate NOT location, around stereotaxic coordinates AP -2.4 to -2.6 mm and ML 1.3-1.4 mm. Duratomy was performed via a 30G needle. The animal was then transferred to the experimental rig for the recording session, where it was placed on a heating pad to keep normothermia (37°-38°C) and its head fixed onto the headpost holder. Oxygen was administered throughout the session. Prior to probe insertion, a drop of low melting point agarose (1.2%) was placed onto the exposed brain to avoid dehydration and for mechanical stability. Vital parameters were monitored every 15 minutes for the whole recording duration via a pulse oximeter and heart rate monitor (MouseSTAT Jr with Paw Sensor, Kent Scientific). A re-dosing scheme was applied dynamically throughout the experimental session based on respiratory rate, i.e., in case the respiratory rate exceeded 150 min^-1^.

### Experimental setup, mice experiments

The experimental setup consisted of an elevated 20 x 10 cm breadboard platform (*Thorlabs*) with a heating pad (*Harvard Apparatus*), placed onto an anti-vibration table and facing one (for neural recordings) or two (for psychophysics measurements) LCD monitors (DELL U2520D). The monitors were placed in front of each eye in a vertical orientation and each spanned 96×126 degrees of visual angle (dva). The platform was clamped on a 37cm-high pillar post, above which a probe micromanipulator (uMp-4, *Sensapex*) was mounted. During the experiment, the anesthetized mouse was placed on the heating pad and its head secured to the headpost holder. The mouse was positioned so that the left eye was pointing at the left monitor’s center, at a distance of 14 cm.

An eye-camera with an infrared (IR) filter (DMK 37AUX287, The Imaging Source) was mounted at the back of the platform and acquired pupil position images from the left eye via a hot mirror placed at the front of the platform during psychophysics recordings. The eye was illuminated by an IR light source placed below the platform, to the left side.

A movable stereomicroscope was placed on a platform, to the side of the recording platform, to allow monitoring of probe insertion.

### Visual stimulation, mice experiments

Stimulus presentation was controlled using custom MATLAB scripts and Psychtoolbox-3(*59, 60*). Stimuli were presented on gamma-corrected displays at a 2560 × 1440 resolution and a refresh rate of 60 Hz.

The stimulus set comprised square-wave drifting gratings (spatial frequency, 0.025 cycles/degree; speed, 45°/s), a full-field Chirp stimulus (CS(*61*)), random moving objects (RMOs(*41*); object size, 20°; speed, 45°/s; object density: ∼5.5 objects were, on average, visible on screen in each frame), binary white noise (pixel size, 10°; 20 Hz refresh rate), and coherently moving square, left-sheared, and right-sheared objects (object size, 20°; speed, 45°/s; object density: ∼20.8 objects were, on average, visible on screen in each frame). Coherently moving squares had the same appearance as RMOs, but all objects moved in a common direction.

Drifting gratings and coherently moving square and sheared objects were presented in eight directions, separated by 45°, with five trials per direction in randomized order. Each trial lasted 12.5 seconds, comprising 10 s of coherent motion followed by a 2.5-second inter-trial interval. For drifting gratings, the interval consisted of a blank (black) screen. For square and sheared objects, it consisted of incoherent object motion (objects moving in random directions). Each trial of square and sheared objects contained an independently generated random sequence of moving objects, such that repeated presentations of the same motion direction were not identical.

The CS was presented at the beginning and end of each recording session, with four or eight repetitions, to identify visually evoked activity and neural longevity between start and finish of the recording session. Cells that did not retain their visually evoked activity by the end of the recording session were excluded from further analysis. RMOs and binary white noise were presented continuously for 20 min, without interleaved inter-trial interval periods.

‘Sheared’ objects were generated by transforming square objects using a two-dimensional shear matrix, where each vertex (x,y) was mapped to (x+ky,y). For a 45° shear, the parameter was set to k=tan(45°)=1, and for the opposite shear, k was set to −1, producing equivalent distortions in opposite directions (Fig. 1D). Stimulus size was calculated as follows.

The visual angle, θ, was determined using the formula

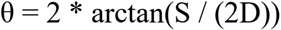

where S represents the size of the stimulus and D denotes the distance from the mouse to the screen. Specifically, S was measured in µm and corresponds to the physical dimensions of the stimulus on the screen, while D was the distance from the mouse’s eyes to the screen (14 cm). The visual angle was then converted into µm on the retina based on an approximated eye size of 3300 µm(*62*) as previously reported(*63*).

### Animal preparation, experimental setup and visual stimulation: marmoset experiments

#### Participants

Two adult common marmosets (*Callithrix jacchus*), Marmoset A (male) and Marmoset M (female) were used for eye tracking experiments with motion stimuli. Subjects were housed at the University of Rochester with a circadian cycle of 12-hour light and 12-hour dark.

#### Preparation

All subjects were surgically implanted with head caps to stabilize them for head-fixed eye tracking as described previously(*64, 65*). Marmosets were then trained to sit in a primate chair under head restraint while facing a projection screen with vision corrected by a lens(*64*).

#### Visual stimulation

Visual stimuli were presented on a Pro Pixx Projector (*VPixx Technologies*) with a linear gamma running at 80 Hz at 1280 x 720 resolution. Stimuli were generated via MATLAB’s Psychophysics ToolBox (*59, 60*). Task events were recorded using a DATAPixx I/O box (*VPixx Technologies*) for temporal registration.

The MATLAB code is available online (https://github.com/jcbyts/MarmoV5) and was used in previous studies(*66*). The stimulus screen was viewed at a distance of 68 cm and subtended ± 24 horizontally and ± 13.5 vertically in dva.

Motion trials sampled between square stimuli spanning 1×1 dva, a square of the same size with a left or a right shear. To prevent a rigid grid like appearance of square or sheared stimuli, each stimulus was placed within a grid of 2 x 2 dva sections with the center displaced at random by ± 0.375 dva horizontally and vertically (uniformly sampled from -0.375 to 0.375 dva). The stimuli drifted at 10, 20 or 40°/s, in interleaved trials, in one of the four cardinal directions (up, right, left, down) for a duration of 10 seconds followed by a 2-3 second inter-trial interval. No qualitative difference was observed in eye movement directions across different speeds. The data shown in Fig. 2G represent eye movements collected at 20°/s. During each trial, marmosets were allowed to free view the stimulus and received a drop of liquid reward (5 micro-liters) at the trial end. Due to the limited oculomotor range of marmosets(*67*), they generally viewed within ±10 dva of the screen center.

### Neural recordings

Neural recordings were performed extracellularly, from the right hemisphere NOT, via Neuropixels(*37*). Before the recording, a silver wire was permanently soldered to the probe ground pin. During the recording, the silver wire was attached to the headpost holder, and an internal reference (the probe tip) configuration was used. The probe was mounted on a dovetail probe holder, secured to the micromanipulator and inserted into the brain at a speed of 2 µm/s until the desired depth, typically around 4-4.2 mm. Stereotaxic coordinates of insertion were usually in the range AP -2.6 ± 0.2, ML 1.4 ± 0.1 and DV 4 ± 1 mm. The probe was left to settle for 15 minutes before recording started, to minimize mechanical drift.

Visually-evoked activity was assessed upon electrode insertion in response to the Chirp stimulus. Direction selectivity to temporo-nasal motion was then assessed by showing drifting gratings. The electrode was repositioned until robust direction selective temporo-nasal responses were identified.

Neural signals were band-pass filtered (300-5000 Hz) and acquired at 30 kHz via the *Openephys* GUI (https://open-ephys.org/) at a gain setting of 300x. Automatic identification of spike times and the assignment of spikes to individual neurons was achieved using the Kilosort2.5 MATLAB package (https://github.com/MouseLand/Kilosort).

Offline spike curation was performed using *Phy* (https://github.com/cortex-lab/phy). Good units were selected based on similarity of waveform shape, spike amplitude, lack of violation of the refractory period, and lack of fluctuations due to mechanical drift. Well-isolated units that stopped their activity during the recording session were marked as ‘noise’ and excluded from further analysis.

### On-line NOT localization

After visually evoked activity was confirmed with a full-field Chirp stimulus, drifting gratings were used to assess direction selectivity, particularly for temporo-nasal motion. To further verify correct targeting of NOT, a 20° × 20° white square moving in the temporo-nasal direction was presented. This stimulus provided a rapid estimate of receptive field size and location, allowing us to distinguish NOT from the neighboring superior colliculus (SC) at the start of the recording: although posterior SC neurons can also exhibit temporo-nasal direction selectivity(*68, 69*), their receptive fields are typically small (∼10°), symmetric, and span the whole visual field depending on probe insertion coordinates(*70*). In contrast, our data show that NOT receptive fields are larger, elongated, and cluster near the horizontal meridian. Once the characteristic NOT functional profile was identified, visual stimulation experiments were initiated.

### Data synchronization

Neural recordings were performed using an acquisition system consisting of a PXIe-1000 chassis (*National Instruments*) and an HST-1000 headstage interface. To synchronize stimulus presentation, eye-tracking data, and neural activity, we employed a separate NI PXIe-1071 4-slot chassis configured with a digital I/O DAQ card.

Three computers were used during the experiments: A1 for visual stimulus presentation, A2 for eye tracking, and A3 for Neuropixels data acquisition (running *Openephys*). TTL pulses were sent from A1 (at each stimulus onset) and from A2 (for behavioral event marking) to the NI PXIe-1071 chassis. These pulses were recorded and timestamped to allow precise alignment of stimulus and behavioral events with the neural data. The PXIe-1071 was connected to the Neuropixels acquisition system, enabling shared timing signals between systems.

### Histology, immunohistochemistry and anatomical assignment of neurons

Post-recording, the Neuropixel probe was extracted, coated with a fluorescent dye (DiI, *Thermo Fisher*), and reinserted to the recording depth. This allowed us to identify the insertion coordinates of the electrode by imaging the fluorescence left by the electrode track. Then, animals were placed under overdosing Isoflurane anesthesia and underwent transcardiac perfusion with 1 × phosphate-buffered saline (PBS), followed by 4% paraformaldehyde (PFA). The brain was then extracted and kept in PFA for 24 hours before being transferred to PBS and left there for 1-7 days. The brain was sliced into 75 µm-thick sections using a vibratome, and the slices were placed in PBS in a 24-well plate. Brain slices of WT mice were mounted directly after this step, and the electrode track visualized via stereomicroscope without immunological enhancement. Brain slices of Hoxd10 mice were stored in multiwells in PBS. PBS was then removed, and samples were incubated in 300 μl blocking solution composed of 10% normal donkey serum (NDS; Sigma-Aldrich, S30-M), 1% BSA (Sigma-Aldrich), 0.02% Sodium Azide (NaN3; Sigma-Aldrich, S2002), 0.5% Triton X-100 (Sigma-Aldrich, 93443) and 1× PBS for 2 h under shaking conditions at room temperature. For antibody incubations, the same buffer was used with 3% NDS. Samples were then incubated for 2 days at room temperature under shaking conditions in primary antibody solution (rabbit anti-GFP polyclonal; *Thermo Fisher Scientific*; catalog no. A-11122, RRID: AB_221569; 1:200). After three PBS washes, the secondary antibody solution (Alexa Fluor 488-conjugated donkey anti-rabbit IgG (H+L), highly cross-adsorbed; *Thermo Fisher Scientific*, catalog no. A-21206, RRID: AB_2535792; 1:200) was applied and left overnight under shaking conditions at room temperature. The following morning, the slices were washed with PBS, placed on glass slides, and mounted with ProLong (*Invitrogen*). Images of the brain slices were acquired using a Spinning Disc Confocal system (*Yokogawa Electric*) on a microscope operated by the *CellSens* Software (*Olympus*). The NOT was identified by the presence of GFP-fluorescent axonal terminals below the cortical layers and anterior to the Superior Colliculus. Coordinates of insertion were marked by the red fluorescent trace left by the DiI-coated electrode. The known geometry of Neuropixels allowed us to triangulate direction-selective channels with the anatomical landmark of NOT (fig. S1).

### Analysis of neural responses

All data analysis was performed via custom-made MATLAB and Python 3.8 routines. Our analysis was based on single units isolated via spike sorting and spike curation. We included neurons that exhibited a firing rate of at least 0.1 Hz over the full duration of each presented stimulus and showed robust light responses to at least one parameter combination of the drifting gratings stimulus, determined as follows: first the neuron fired at least 30 spikes per trial throughout the stimulus duration; second, the absolute firing rate modulation index exceeded 0.5; third, the maximum trial-to-trial correlation coefficient (see below), with the maximum taken over the grating directions, exceeded 0.3.

#### Firing rate estimation

Scalar firing rates of neurons during stimulus presentation were calculated by dividing the number of spikes during the stimulus presentation by the duration of the stimulus presentation. Time-resolved firing rates for each trial were calculated via kernel density estimation(*71*) using a Gaussian kernel (Δt=10ms, *σ* = 50ms) and then averaged over trials.

#### Trial-to-trial correlation coefficient

Reliability of neural responses was quantified using a trial-to-trial correlation method described elsewhere(*72*). In short, spike trains were binned in time (Δt = 250 ms) to form a peri-stimulus time histogram. Binned spike counts were then correlated between all possible combinations at once. For example, in the case of *n*_*trial*_ = 3, three binned spike count vectors were obtained *a*_1_, *a*_2_, *a*_3_ and arranged into two vectors *v*_1_, *v*_2_ by concatenation as follows:

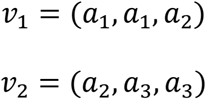

The trial-to-trial Pearson correlation coefficient *ρ* was computed as well as its statistical significance using MATLAB’s *corrcoef* function.

#### Absolute firing rate modulation index

As an estimator for the quality of visually-evoked neural activity, we computed a firing rate modulation index based on a neuron’s response to the drifting gratings (MG) and CS stimuli separately. The MG firing rate modulation index was defined as follows:

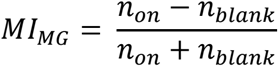

where *n*_*on*_ was the spike count during the stimulus period of the MG summed across trials, and *n*_*blank*_ was the spike count during the blank period of the MG summed across trials.

The CS firing rate modulation index was defined as follows:

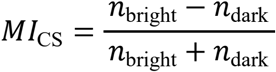

where *n*_bright_ was the spike count in a 1s interval after the start of the bright contrast step and *n*_dark_ was the spike count in a 1s interval at the beginning of the CS trial, corresponding to a dark background. Cells were determined to be light responsive to the CS stimulus when they exhibited a significant, positive trial-to-trial correlation coefficient (ρ > 0, p < 0.05) and the absolute firing rate modulation index |*MI*_CS_| exceeded 0.25.

#### Direction selectivity index

A direction-selectivity index (DSI) was computed to determine DS-RGCs in the mouse retina (in response to a moving bar stimulus) and NOT (in response to drifting gratings stimuli). Spike counts were computed for each trial and 8 directions. Tuning curves *c*(*α*) ∈ *R*^(8×1)^ as a function of direction *α* were computed by taking the median spike count across trials. A DSI(*61*) was then computed by projecting the tuning curve on a complex exponential *ϕ*_*k*_ = *e*^(*iαk*)^, where α_k_ was the k-th direction:

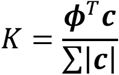

The DSI was then defined as the magnitude of the complex number K:

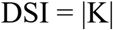

Significance of the DSI was determined by a permutation test. For each cell, direction-dependent responses were arranged in a direction-trial response matrix. Entries in this direction-trial matrix were randomly permuted (both over trials and over directions), destroying any relationship between the stimulus and the response, before computing the permuted DSI. We repeated the process 1,000 times to create a null distribution for the DSI. The p-value was the fraction of permuted DSIs exceeding the true DSI. A DSI with p < 0.05 was determined as significant. A cell was determined as direction selective when DSI ≥ 0.2 and p < 0.05.

#### Receptive field estimation with random moving objects

Receptive fields were estimated from responses to random moving object stimuli (RMO) using the Reverse Correlation Against Stimulus Elements (RCASE) framework(*41*). A stimulus element, i.e. an object in the RMO stimulus is described in terms of its parameters (position, size, speed, direction). The edge size of an object covered 20° of visual angle.

Object starting positions were sampled uniformly across the screen (x,y), movement directions sampled uniformly between 0° and 360°. The time of appearance of each object was sampled uniformly across the total stimulus duration and objects remained on the screen for up to 3 seconds before disappearing. To construct the stimulus design matrix, stimulus elements parameters were discretized: object center positions were binned into square spatial bins of 200μm on the retina (∼7° of visual angle), while motion directions were binned in 45° steps.

Each object is encoded using one-hot vectors for its parameters, which are combined via a tensor (outer) product to form a high-dimensional binary tensor with a single non-zero entry. Within each frame, single object tensors are summed across objects. The resulting summed tensors per frame are concatenated across time and vectorized to form a sparse stimulus design matrix *X* ∈ ℕ^*T*×*D*^, where *T*is the number of frames and *D* the stimulus dimensionality.

Neural responses were binned at the same temporal resolution (Δ*t* = 1/60 s) to obtain a spike count vector *y*.

Receptive fields were computed by reverse correlation between *X*and temporally shifted responses *y*_*τ*_, which can be expressed in terms of a linear regression problem:

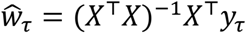

where *τ* denotes the time lag. Spatiotemporal receptive fields *ŵ* are obtained by evaluating *ŵ*_*τ*_over lags in the range [−2 s, 2 s]. For efficiency, the (*X*^⊤^*X*)^−1^*X*^⊤^ is computed once and reused across lags.

While this stimulus has no spatial correlations, because objects persist across frames, the stimulus exhibits temporal correlations, which were not corrected and should be taken into account when interpreting receptive field estimates.

#### Receptive field estimation with white noise

The white noise stimulus consisted of 10°-wide squares arranged in a checkerboard pattern, changing patterns at a rate of 20 Hz. The stimulus design matrix *X* encoded the contrast of the patterns across time. Each row *X*_*i*,:_ of *X*, represented the vectorized 2-dimensional white noise stimulus of frame *i*. The values of *X* were chosen uniformly at random from {-1,1}, which represented black or white contrast, respectively. Identically to the RCASE analysis, we represented neural responses by vectors of spike counts *y* at the WN frame rate with the same number of elements *T* as rows in *X*. Receptive fields for time lag *τ* were estimated using the spike-triggered average (STA)(*73*):

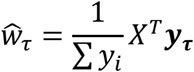

*ŵ*_*τ*_ was the receptive field for a given time lag *τ*, and *y*_*τ*_ was the neural response shifted by a lag of *τ* frames. In total 15 time lags were included to compute the full spatio-temporal receptive field.

#### Decomposition of spatiotemporal receptive fields into spatial and temporal components

We rearranged the 3-dimensional spatiotemporal (*x, y, τ*) receptive fields into two-dimensional matrices, where rows denoted the spatial dimension, and columns denoted the temporal dimension:

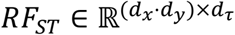

We then applied principal component analysis (PCA) to the spatiotemporal receptive fields and projected the spatiotemporal receptive field on the first principal component. The obtained vector was then arranged in a two-dimensional matrix with the original spatial dimensions, which we defined as the spatial receptive field:

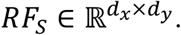

We determined the location of the receptive field center as the coordinates of the pixel in the spatial receptive field with the largest absolute value.

#### Receptive field quality index, size and aspect ratio

We quantified the receptive field quality utilizing a quality index (Qi)(*61*). A two-dimensional Gaussian was fitted to the absolute spatial receptive field, using Matlab’s *lsqcurvefit*. The receptive field quality was defined as one minus the fraction of variance unexplained by the Gaussian fit 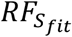:

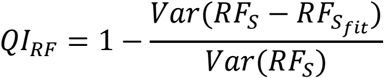

A cell exhibited a receptive field if the Qi exceeded a value of 0.3. The receptive field size was calculated as the mean between the width (2*σ*) of the x and y axes of the 2D Gaussian fit, multiplied by spatial resolution of the stimulus. Aspect ratio was calculated as the ratio between major and minor axis of the 2D Gaussian fit; an aspect ratio of 1 is a circular receptive field, while an elongated aspect ratio exceeded a threshold of 1.3.

#### NOT retinotopy maps

Direction-selective (DS) NOT neurons were assigned retinotopic coordinates by mapping each unit’s receptive-field center onto the mediolateral-anteroposterior (ML-AP) coordinate system defined by the electrode insertion tracks. For each session, the stereotaxic ML and AP coordinates of the probe entry point were propagated to all visually responsive units recorded along that penetration. Receptive field centers were estimated from reverse-correlation receptive-field maps, and their azimuth and elevation coordinates were extracted from the corresponding spatial location of peak response.

To visualize population-level retinotopy, receptive field positions from n=137 DS cells were pooled across experiments and projected into a common ML-AP coordinate space. Two-dimensional retinotopic maps were obtained by interpolating receptive field elevation or azimuth values using a natural-neighbor scattered interpolant (“natural” method in *scatteredInterpolant*; MATLAB, MathWorks), evaluated on a dense 100×100 grid spanning the sampled ML-AP range. To prevent artificial extrapolation outside the empirically sampled region, a masking procedure was applied: for each grid point, the Euclidean distance to the nearest recorded cell was computed, and grid points farther than 0.2 mm from any sample were excluded from the visualization.

Because uneven anatomical sampling could influence the interpolated maps, we quantified the local sampling density by computing a 2D histogram of DS-cell receptive field positions (20×20 bins) in ML-AP space. This receptive field-density map was visualized separately and used to identify sparsely sampled regions in which interpolation is less reliable. For display purposes, density values were normalized to their maximum and plotted either as a standalone heatmap or overlaid on retinotopy maps using contour lines with spatially varying alpha transparency. Scatter plots of individual receptive field centers, colored according to local sampling density, were added to all retinotopic projections to facilitate interpretation of local features.

Together, these procedures generated smooth, interpretable 2D maps of receptive field elevation, receptive field azimuth, and receptive field sampling density across the anatomically sampled portion of mouse NOT, while minimizing artifacts arising from nonuniform penetration coverage.

### Two-stage retina-NOT model

#### Spatial lattice

Responses of NOT neurons to moving objects were modeled as a rectified linear pooling of local edge motion detectors (hereafter, detectors) tiled across the visual field. Local motion signals were computed using a dense array of detectors covering the stimulus display (1440 × 2560 px). Detector centers were arranged on a hexagonal lattice with a spacing of 195 px, alternate rows being offset by half that spacing. A lattice position was kept if the corresponding receptive field, out to one standard deviation from its center (see below), overlapped the display (136 positions). Each detector *j* was defined by a circular Gaussian receptive field centered at (x_j_, y_j_) with standard deviation *σ*_RF_.

The spatial receptive field of detector *j* was defined as:

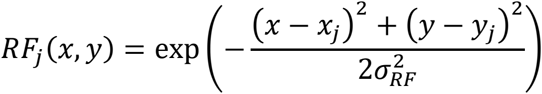

with *σ*_RF_ = 100px.

#### Edge-based local motion computation

For each stimulus frame, object geometry and kinematics were obtained from the stimulus metadata. Objects could be square or sheared (see *Visual stimulation)*. For each object, the four edge segments were reconstructed in screen coordinates, and the leading and trailing edges were identified relative to the instantaneous object motion direction.

Let v_obj_ denote the global object velocity vector. For a given edge with unit orientation vector *e*, the unit normal vector was defined as:

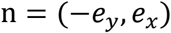

Local edge motion was defined as the projection of the global velocity onto the edge normal (i.e., the aperture-limited normal-flow component):

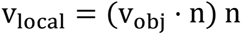

The corresponding local motion direction in screen coordinates was:

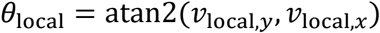

Thus, each edge contributed a local motion signal corresponding to the normal-flow component of object motion.

For each edge, 100 equally spaced points were sampled along the edge segment.

#### Spatial integration

For detector *j*, and edge sample point *i*at position (*x*_*i*_, *y*_*i*_), the spatial weighting was:

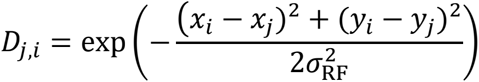

The spatial response of the entire edge was defined as the average across sampled points:

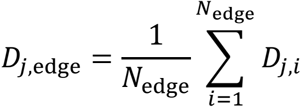

Objects whose center lay farther than 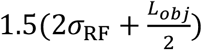 from the center of a detector were not evaluated for that detector, where *L*_*obj*_ is the side length of the unsheared object.

#### Direction selectivity

Direction selectivity was implemented using a normalized von Mises tuning function:

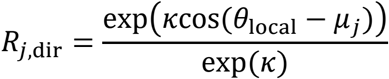

normalized to a peak of 1, where *μ*_*j*_is the preferred direction of detector *j*, and 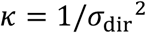 the concentration parameter with a tuning width of *σ*_dir_ = 60^∘^ (in radians). For non-direction-selective detectors, *R*_*j*,dir_ = 1.

#### Frame-wise detector response

For each edge in frame *t*, the detector response was defined as:

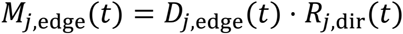

Each detector responded to one edge type, either the leading or the trailing edge. If several edges of that type were present in the same frame, their responses were summed:

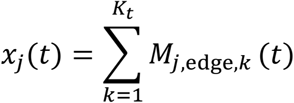

where K_t_ is the number of edges of the detector’s type in frame t. This produced one scalar feature value per detector per frame.

#### Grid model of NOT cells

To simulate NOT responses, three detector grids were constructed over the same hexagonal lattice, each instantiated once for the leading and once for the trailing edge:

1. Direction-selective grid with preferred direction 22.5°
2. Direction-selective grid with preferred direction 202.5° (opponent direction)
3. Non-direction-selective grid

Detector parameters (*σ*_RF_, *σ*_dir_) were fixed across detectors and stimulus conditions. The outputs of all detectors were concatenated into a design matrix:

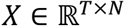

where *T* is the number of frames (36,000; 600 s at 60 Hz) and *N* the number of detectors (816; 136 lattice positions × 3 grids × 2 edges).

#### Divisive normalization

The output of each detector was divided by the non-direction-selective motion signal in its neighborhood, *s*_*j*_(*t*):

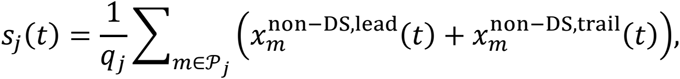

where *P*_*j*_ is the set comprising the lattice position of detector *j* and its nearest neighbors, 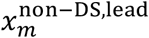 and 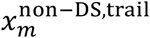 are the outputs of the non-direction-selective detectors for the leading and the trailing edge at position *m*, and *q*_*j*_ is the 99th percentile, over all frames of the stimulus condition, of the summed output in parentheses, so that *s*_*j*_(*t*) ≤ 1 in 99% of frames. The normalized output of detector *j* was

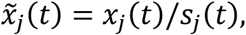

With 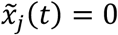 where 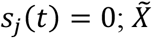 denotes the design matrix of the normalized outputs.

#### Linear pooling model

For each neuron, the predicted firing rate was modeled as a linear combination of the divisively normalized detector outputs plus an intercept, followed by a rectifying nonlinearity:

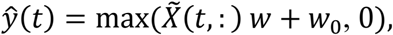

where w is a vector of regression weights (one weight per detector) and *w*_0_ is the intercept. Weights and intercept were estimated by L1-penalized least squares using MATLAB’s *lasso* function, which solves:

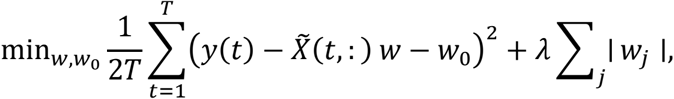

where *y*(*t*) is the neuron’s measured firing rate in frame *t* (see *Measured response and temporal alignment*), and λ set the strength of the penalty (see *Cross-validation performance metrics*). The rectification was not part of the fit: weights and intercept were estimated for the linear stage,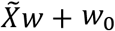, and the rectification was applied to every model prediction that was evaluated. Fits were performed separately for each stimulus condition (‘square’, ‘left sheared’ and ‘right sheared’ objects).

#### Measured response and temporal alignment

The model was fitted to NOT neurons that met the inclusion criteria (see *Analysis of neural responses*) and fired at a mean rate of at least 3 Hz in each of the three stimulus conditions (n = 21 DS and 26 non-DS neurons, the same neurons in all conditions). For each neuron, spikes were counted in bins of one stimulus frame (∼16.7 ms). To reduce response noise, the spike counts were smoothed using MATLAB’s *smoothdata* function with a Gaussian kernel (standard deviation 4 frames, ∼67ms), resulting in the measured firing rate *y*(*t*) to which the model was fitted. To account for neural response latency, the design matrix X was delayed by 6 frames (100 ms) relative to the measured firing rate, the first 6 frames were set to zero.

#### Cross-validation performance metrics

Model performance was evaluated within each stimulus condition using cross-validation on five contiguous blocks, each block containing one stimulus repetition (each of the eight motion directions once).

For each fold the penalty λ was chosen by an inner cross-validation on the four training blocks only. For each neuron, eight candidate values were spaced logarithmically from 10^−4^*λ*_max_ to 10^−0.5^*λ*_max_, where *λ*_max_ was the smallest penalty at which all weights were zero, computed from the training blocks. Each training block was left out in turn, the model was fitted on the other three for each candidate, its rectified prediction of the left-out block was evaluated, and the candidate with the highest R^2^ (defined below) on the left-out blocks, averaged over the four fits, was selected. After selecting λ weights were then fitted on the four training blocks, and the rectified model predicted the held-out block.

Model fit was quantified by the coefficient of determination:

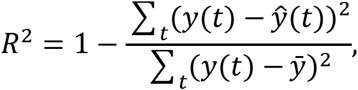

where the sums run over the evaluated frames and 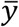 is the mean of *y* over these frames. Model performance was the (outer) cross-validated 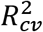 : the predictions for the five held-out blocks were concatenated into one prediction of the whole trace, 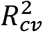, and *R*^2^ was computed once over all 36,000 frames. For population summaries, the median *R*^2^ across neurons was reported. For analyses of the model weights (Fig. 5B-C; fig. S11, A, D-E) the weights of one fit to all frames at the median of the five selected λ values were used.

#### Tuning curve analysis

Measured and predicted tuning curves were computed by summing the smoothed measured firing rate and the cross-validated prediction within periods of coherent object motion. For each direction the median was taken across trials with the same direction of motion. For the population tuning of DS neurons, the measured and predicted curves of each neuron were divided by the maximum of its measured curve and averaged across neurons. Agreement between measured and predicted population tuning was quantified by the *R*^2^, computed between the mean measured and the mean predicted curve over the eight directions.

#### Receptive field/weights grid correlation analysis

The dominant grid of a neuron (Fig. 5D) was defined as the grid with the largest summed absolute weight (leading and trailing edge). Spatial correspondence between model-derived weights and receptive fields was quantified for neurons with a receptive-field fit quality index (Qi) > 0.3. A continuous model-derived spatial map was constructed by placing the Gaussian receptive field of each lattice position at that position. Each Gaussian was scaled by the summed absolute weights of all detectors at that position (three grids, two edges), and the scaled Gaussians were summed. The map was averaged within each spatial bin of the receptive-field estimate (33 × 19 bins). Spatial correspondence was quantified using Pearson correlation between the model-derived map and the two-dimensional Gaussian fit of the neuron’s receptive field across all bins.

#### Model comparison

The divisive normalization model was compared to a linear model fitted with maximum likelihood estimation (Ordinary Least Squares) on non-normalized inputs, a linear model fitted with L1-Regularization on non-normalized inputs, and a nonlinear boosted regression tree model fitted on non-normalized inputs. Unless stated otherwise, the comparison models used the same subunits, target *y*(*t*), cross-validation blocks and rectifying nonlinearity as the divisive normalization model, and were evaluated by the same 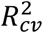 .

#### Model comparison – Maximum Likelihood

The weights of the non-normalized subunit outputs *x*_*j*_ and the intercept were estimated by ordinary least squares without penalty, solved from the normal equation

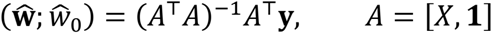

where *X* is the matrix of the non-normalized subunit outputs *x*_*j*_(*t*) and **y** the response vector across frames. As the model has no hyperparameter, each held-out block was predicted by the fit to the four training blocks, and the weights shown in the maps came from one fit to all frames.

#### Model comparison – Lasso

The weights were fitted with the lasso objective on the non-normalized subunit outputs *x*_*j*_(*t*) as for the divisive normalization model above with the objective:

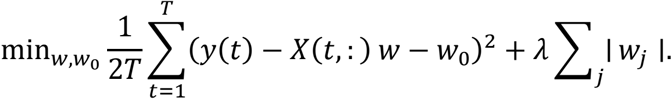

#### Model comparison – Boosted trees

The response of each neuron was modeled as a sum of regression trees on the non-normalized subunit outputs *x*_*j*_(*t*). Trees were fitted one after the other, each to the residual of the trees before it (boosting), their predictions were scaled by the learning rate and summed, and the sum was rectified as in the other models. The ensemble was fitted with MATLAB’s *fitrensemble* (method ‘LSBoost’), with the default tree template *templateTree* as the weak learner. All hyperparameters were kept at their MATLAB defaults, except for three: the learning rate (*LearnRate*) was set to 0.1, each input was binned into at most 50 equiprobable bins (*numBins*), and the number of trees. The number of trees, tested from 1 to 500, was the only fitted hyperparameter and was chosen by the same cross-validation procedure as *λ* in the lasso or divisive normalization model. In place of weights, to compute weight maps and calculate the RF overlap, the feature importance of each subunit was computed from one ensemble fitted to all frames with the median of the five selected numbers of trees using MATLAB’s *predictorImportance*.

### HDMEA recordings

The high-density microelectrode array (HD-MEA) recordings were performed on a CMOS-based chip featuring 26,400 electrodes with a center-to-center pitch of 17.5 μm, and 1024 parallel readout channels(*47*). Each readout channel featured an analog-to-digital converter (ADC) with 10-bit precision and 20 kHz frame rate.

The explanted retina from the enucleated eye was separated from the retinal pigment epithelium and placed ganglion-cell-side down on a HD-MEA. Using a micromanipulator, the retina was pressed onto the electrodes with a transparent polyester tissue-culture membrane (10 μm thick, 0.4 μm pore size, product number 3450, Corning, New York, United States) in order to increase the tissue attachment to the MEA and to keep the tissue in place during the recording.

The membrane was modified to increase tissue perfusion with holes at regular spacing (100 μm radius in a 400 μm raster, IMPEX Leiterplatten GmbH, Michael im Lungau, Austria). The temperature was kept at 38°C with the help of an active perfusion of 2.5 ml/min and an inflow temperature controller.

For each retina piece, placed on the array, we first mapped the activity across the entire array to locate areas best suited for the experiment. We then selected a high-density electrode configuration in the selected region according to design constraints of the switch-matrix.

Once a region was selected, we performed gray adaptation by exposing the cells to one minute of gray screen. After this time, light stimuli were shown centered onto the middle of the recording area covering all selected electrodes. All stimuli were generated on MATLAB’s Psychtoolbox(*59, 60*) and had the same parameters used for NOT experiments, except for

- The use of a 20° full field moving bar, instead of drifting gratings, moving at 10°/s and 45°/s, for direction-selectivity assessment for direction-selectivity assessment.
- The size of the RMO stimulus for receptive field mapping (7°), with spatial bins of 3.5°. Spike sorting and curation of HD-MEA data were performed as described in(*74*).

### Mouse psychophysics

#### Eye movements acquisition and calibration

Eye-movement videos were acquired from the left eye using an infrared camera positioned behind the animal. The camera viewed the eye through a hot mirror that reflected infrared light while remaining transparent to the visual stimulus (fig. S3, B; fig. S4, B). Pupil position was extracted offline using custom MATLAB code. Video frames were converted to grayscale, contrast-enhanced, and thresholded to isolate dark pixels corresponding to the pupil. A manually selected polygonal region of interest (ROI) was applied across frames, and the largest connected dark component within the ROI was identified as the pupil. Its centroid defined the raw two-dimensional pupil position,

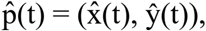

expressed in pixel coordinates. Pupil size was estimated from the average of the major and minor axis lengths. Tracking quality was visually inspected using videos with the detected pupil overlaid on each frame.

Pupil displacements were converted into degrees of eye rotation using an eye-specific calibration based on two infrared LEDs rigidly mounted on the camera housing (fig. S3, B-C). The LEDs were separated by a known distance (d_1_ = 1.5 cm) and positioned at a distance d_2_ from the center of the mouse eye, subtending a visual angle:

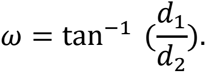

The reflections of the two LEDs on the eye surface were manually identified from a representative video frame, and their separation (D_i_) was calculated as the Euclidean distance between their image coordinates. Assuming approximately spherical eye geometry, the calibration related the subtended angle to twice the measured reflection separation. The Hirschberg ratio (*75, 76*), i.e. the conversion factor between pupil displacement in pixels and eye rotation in degrees, was then calculated as:

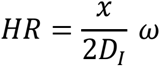

where x is the pixel-to-millimeter calibration factor obtained from an image of a ruler placed in the focal plane of the eye. The resulting conversion factor was applied to pupil displacements to obtain calibrated eye-position traces,

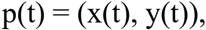

expressed in degrees of eye rotation.

OKR events were identified manually from the calibrated eye-position traces as pairs of slow (ramp) and fast (reset) phases (fig. S3A). Unless otherwise specified, the beginning (t_1_) and end (t_2_) of each slow phase were selected from the horizontal (yaw) eye-position trace. For comparisons of horizontal and vertical OKR event numbers (fig. S4A), events were identified separately from the horizontal (yaw) and vertical (pitch) traces. Manual identification was not blinded to stimulus condition but was based on the characteristic slow and fast phases, independently of subsequent quantitative analyses.

For each OKR event, the two-dimensional eye-displacement vector was calculated between the beginning and end of the slow phase:

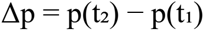

#### Eye movements direction

Eye-movement direction was determined relative to an external reference axis defined by a separate pair of infrared LEDs mounted on the stimulus monitor and imaged on the eye surface (fig. S4B). The LED coordinates defined a reference vector (r) and its midpoint, which served as the reference center (fig. S4C). Pupil positions were re-centered and reflected across the horizontal axis to match visual-field coordinates. The signed angle between the eye-displacement vector (Δp) and the LED reference axis was calculated using their dot and cross products:

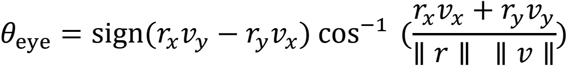

yielding the direction of each eye movement in degrees relative to the horizontal reference axis.

### OKR gain and number of events

OKR responses were measured using 0.025 cpd drifting gratings and 20° square and sheared objects moving at 45°/s, matching the speed used for receptive field estimation. Although horizontal OKR gain decreases with increasing stimulus speed (*38, 77–79*) the higher speed increased the frequency of object passages through individual receptive fields, facilitating the detection of multiple horizontal OKR events.

For each OKR event, horizontal eye velocity was calculated from the horizontal component of pupil displacement over the duration of the slow phase:

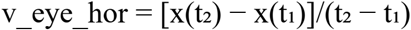

Horizontal OKR gain was defined as the ratio between horizontal eye velocity and stimulus speed:

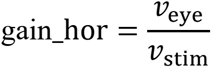

For stimuli containing oblique motion components (including obliquely drifting gratings and square objects, as well as sheared objects irrespective of their global motion direction) horizontal OKR gain was normalized using a fixed 45° projection of stimulus velocity. Specifically, stimulus speed was multiplied by 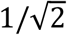, corresponding to the horizontal component of a velocity vector oriented at 45° relative to the horizontal axis:

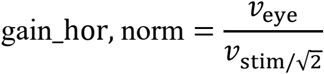

This factor corresponds to the horizontal projection of a velocity vector oriented at 45° relative to the horizontal axis. For gratings and square objects moving along cardinal directions, the full stimulus speed was used.

#### OKR gain symmetry axis

The OKR symmetry axis was quantified to test whether gain values differed across stimulus conditions (Fig. 2A). For each stimulus direction (eight directions spaced at 45°), the median absolute horizontal OKR gain was calculated from trials pooled across mice. Gains for opposite directions (e.g., 0° and 180°) were averaged and assigned to both directions, yielding an axial representation of gain tuning. The symmetry axis was then estimated using the second-harmonic circular moment:

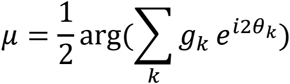

where *g*_*k*_is the symmetrized gain and *θ*_*k*_the stimulus direction in radians.

This estimator identifies the principal orientation of the OKR gain distribution by treating opposite motion directions as equivalent.

### Marmoset psychophysics

#### Eye tracking and calibration

Eye position was recorded using a custom digital Dual-Purkinje Imaging system (dDPI). The dDPI used a collimated IR beam (*ThorLabs*), a dichroic mirror (Edmunds) with the eye imaged on axis, and sampled 0.4 Megapixel images of the eye at 539 frames per second (DMK33UX287; The Imaging Source). Custom CUDA code running on a gaming GPU (GTX1080Ti; *Nvidia*) extracted the gaze position based on the 1^st^ and 4^th^ Purkinje images as described previously(*66, 80*). Eye position was calibrated at the start of each behavior session by showing head-fixed marmosets sets of Gaussian windowed face stimuli as described previously(*64, 67*). Once a rough calibration was obtained by registering x and y eye position to overlap fixations of face stimuli, then it was fine-tuned by running a fixation task with a small point (0.3 dva radius) at the screen center with liquid reward given for holding on the point(*67*).

Eye position was collected during the entire recording session and smoothed offline with a Gaussian kernel (*σ* = 5 ms) as described in a previous study(*81*). Blinks and any epochs in which the registered eye position fell beyond ±15 dva were excluded from analyses.

#### OKR events detection

Eye-position data were analyzed offline using custom MATLAB scripts. Trials were classified according to stimulus type (‘square’, ‘left sheared’, or ‘right sheared’ objects) and global motion direction (upward, downward, leftward, or rightward). Before analysis, trials were subjected to a data-quality criterion requiring valid, finite eye-position measurements in both the horizontal (yaw) and vertical (pitch) components for at least 50% of the trial duration. Trials failing the corresponding threshold were excluded. Missing-data intervals were preserved without interpolation, and no additional smoothing was applied to the previously smoothed eye-position signals.

Similarly for the mice eye movements data, optokinetic events were identified manually from continuous eye-tracking recordings from contiguous low-velocity intersaccadic eye movement segments.

For each OKR event, eye movement trajectories were calculated from the start-to-end displacement vector using the horizontal and vertical eye position components. For horizontal-motion experiments, peaks and troughs were selected from the yaw eye-position trace; for vertical-motion experiments, peaks and troughs were selected from the pitch trace.

The orientation of each trajectory was calculated as:

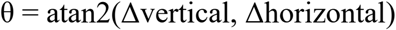

yielding a signed angle representing eye-movement direction in the stimulus coordinate frame.

### Human psychophysics

#### Participants and visual stimulation

Human psychophysics was performed in 10 healthy participants (7 male, 3 female; age range: 29-43), reporting normal or corrected-to-normal vision. Visual acuity values are reported in Table S8. Visual stimuli were generated offline using custom MATLAB scripts (MathWorks) and presented in virtual reality using *SightLab* (*WorldViz*, https://www.worldviz.com) running on the Vizard 8 platform. Stimuli consisted of drifting gratings, square objects and ‘left’ and ‘right sheared’ objects (size: 5°; speed: 45°/s), presented on an extended virtual canvas positioned 1.2 m in front of the participant and moving along the cardinal directions. Each movie consisted of a 5 s blank period, followed by 30 s motion in one direction, 5 s blank, 30 s motion in the opposite direction, and a final 5 s blank period. Subjects were instructed to follow the pattern motion, and avoid focusing on single objects or bars, to curb smooth pursuit behavior. When presented with a central gaze mask, a black, 33° diameter circle, with a 2° Gaussian outer fade, was applied at the center of the visual field, to ensure coverage of the foveal and parafoveal visual field. Subjects were instructed to keep their gaze inside of the confines of the mask, preventing central vision, and only eye movement trajectories falling inside the confines of the mask were included in the analysis.

#### Eye tracking and calibration

Eye movements were recorded using the integrated eye-tracking system of the Meta Quest Pro (https://www.meta.com/ch/en/quest/) headset. Calibration was performed at the beginning of each session using the device’s built-in calibration routine, in which participants followed a moving fixation target across the visual field.

*SightLab* exports eye position in the yaw and pitch axis for each eye at each time point. For consistency with the mouse data, only the signal from the left eye was used for analysis. Eye position values are expressed as Euler angles in degrees, and combined with the headset pose to estimate gaze direction relative to the stimulus plane.

Horizontal and vertical eye-position components were extracted from the eye positions in the yaw and pitch axes, respectively. In contrast to mice, no external calibration reference (e.g., LED-based axis) was required, as gaze direction was directly estimated in world coordinates by the headset.

#### OKR events detection

Eye-position traces were analyzed offline using custom MATLAB scripts. Each trial was divided into two segments corresponding to the two motion directions (leftward/rightward or upward/downward) using a fixed temporal split (35 s).

Similarly for the mice and marmosets eye movements data, optokinetic events were identified manually from continuous eye-tracking recordings from contiguous low-velocity intersaccadic eye movement segments.

### Phenomenological gain model

OKR eye movements were modeled using a two-parameter phenomenological model relating global and local stimulus motion to predicted eye-movement direction and gain. Object motion directions spanned −180° to 180° in 10° steps. For square stimuli, local and global motion directions were identical. For ‘left’ and ‘right’ sheared stimuli, local motion directions were shifted relative to global motion by +45° and −45°, respectively. The contribution of this local-motion shift was controlled by a *bias-strength* parameter ranging from 0 to 1: a value of 0 corresponded to eye movements driven entirely by global object motion, whereas a value of 1 corresponded to the full ±45° local-motion shift. Stimulus motion direction was represented by a two-dimensional vector whose horizontal and vertical components were weighted independently to compute the effective eye motion direction. Their relative weighting was described by a *gain ratio*, defined as the ratio of vertical to horizontal gain. In the isotropic model, horizontal and vertical gains were equal (gain ratio = 1). In the anisotropic model, horizontal gain exceeded vertical gain.

For the mouse model, horizontal and vertical gains were set to values representative of the experimentally observed gain distributions, corresponding to a gain ratio of 0.08, and bias strength was set to 1. For the human model, horizontal and vertical gains were set equal and bias strength was set to 0.2. These parameter values were chosen by hand to illustrate the behavioral regimes observed in the two species and were not obtained by fitting or optimization.

Predicted eye-movement directions were calculated from the weighted horizontal and vertical vector components of the local and global stimulus directions. Predicted gains were calculated as the vector length of the resulting vectors. Model predictions were compared with measured responses to square, ‘left sheared’, and ‘right sheared’ stimuli. For each mouse, we averaged over trials the direction and gain independently for each stimulus type and global motion direction. Angular separations between responses to ‘left’ and ‘right sheared’ stimuli were calculated as signed circular differences.

### Statistics

All statistical analysis was performed in MATLAB.

Direction selectivity indexes (fig. S2, C) across stimulus conditions were compared via a paired permutation test on medians with Bonferroni correction. Differences in direction tuning shifts between conditions were calculated for each cell as the circular difference between paired preferred directions, wrapped to the range -180° to 180°. Statistical significance was assessed using two-sided paired sign-flip permutation tests on the median shift, with Monte Carlo and Bonferroni correction applied across the six pairwise comparisons (Fig. 1G, Table S1).

Differences in the OKR symmetry axis (Fig. 2A) between drifting gratings and moving objects were assessed via a bootstrap comparison on medians.

For each mouse and stimulus condition, OKR gain was calculated by first averaging the gains of individual events within each trial, and then taking the median across trials; the number of OKR events was calculated by counting the detected events within each trial and then averaging these counts across trials. Paired statistical comparisons were performed on these mouse-level values using Wilcoxon signed-rank tests. For visualization only, trial-level values were normalized to common reference values across the conditions being compared: the maximum signed trial-level gain for gain, and the maximum mouse-level mean event count for the number of events. Statistical tests were performed on unnormalized values calculated for each mouse and stimulus condition. Matched conditions were compared using paired Wilcoxon signed-rank tests. For a single prespecified comparison, no multiple-comparison correction was applied, and the uncorrected P value is reported (Fig. 2D-E). For multiple pairwise comparisons across stimuli and motion directions, P values were adjusted using the Benjamini-Hochberg false-discovery-rate procedure (Fig. 4E, center and right panels; Fig. 2B-C).

Direction and magnitude of the angular separation in eye movement direction (Fig. 2F-I; fig. S6) were assessed via a linear mixed-effects model (LMM) for repeated measures, with sheared condition as fixed effect and subject as random intercept. Analyses were performed separately for rightward- and upward-moving objects and for mice, marmosets, humans tested with a central gaze mask, and humans tested without it. The effect of shear condition was quantified as the model-estimated difference between the ‘left’ and ‘right sheared’ conditions, together with its 95% confidence interval and P value. In humans, the effect of masking was additionally assessed by directly comparing the shear-condition effect between masked and unmasked viewing conditions with an LMM with interaction.

Firing rates across stimulus conditions were compared via a paired permutation test on medians with Bonferroni correction (Fig. 4E, left panel; Table S7). Quality indexes (Qis) of receptive fields estimated with different methods were compared via a non-corrected permutation test on medians (fig. S8B).

Tuning curve widths and receptive field metrics in the RGCs population (fig. S10, D-E) were compared via a Ranksum test on medians.

For all bootstrap analyses and permutation tests the number of iterations was 10000, unless otherwise specified.

All tests are specified in the figure captions.

P-values and relevant metrics not reported in the text or figure captions are reported in Tables S1 to S7.

No statistical method was used to predetermine sample sizes.

**Fig. S1.**
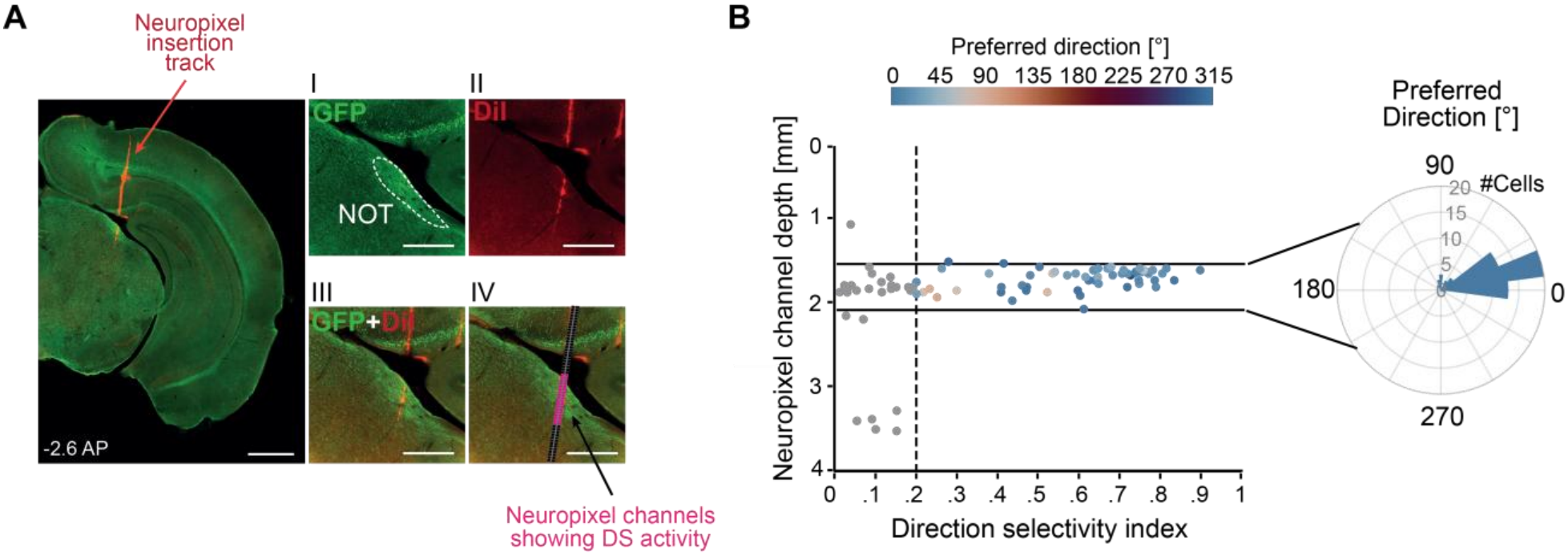
Post-recording localization of NOT in Hoxd10 mice. **A**, Left panel: labeling of a coronal brain slice of an Hoxd10 mouse, right hemisphere, showing the coordinates of electrode insertion (red track). Scalebar: 1mm. Right panel: magnified cut out of the brain slice showing NOT in different color channels; I: GFP labeled T-On-RGCs axonal terminals; II: DiI fluorescent electrode track; III: GFP-labeled NOT with electrode track overlaid; IV: same image in III with overlaid schematics of Neuropixel channels showing a DS response during recording. Scalebar: 500 µm. **B**, Left panel: preferred direction of n=104 NOT cells plotted as a function of direction selectivity index and electrode channel depth. DS cells are located at the depth identified from the histological reconstruction in **A**. Right: distribution of preferred directions for the same cells.

**Fig. S2.**
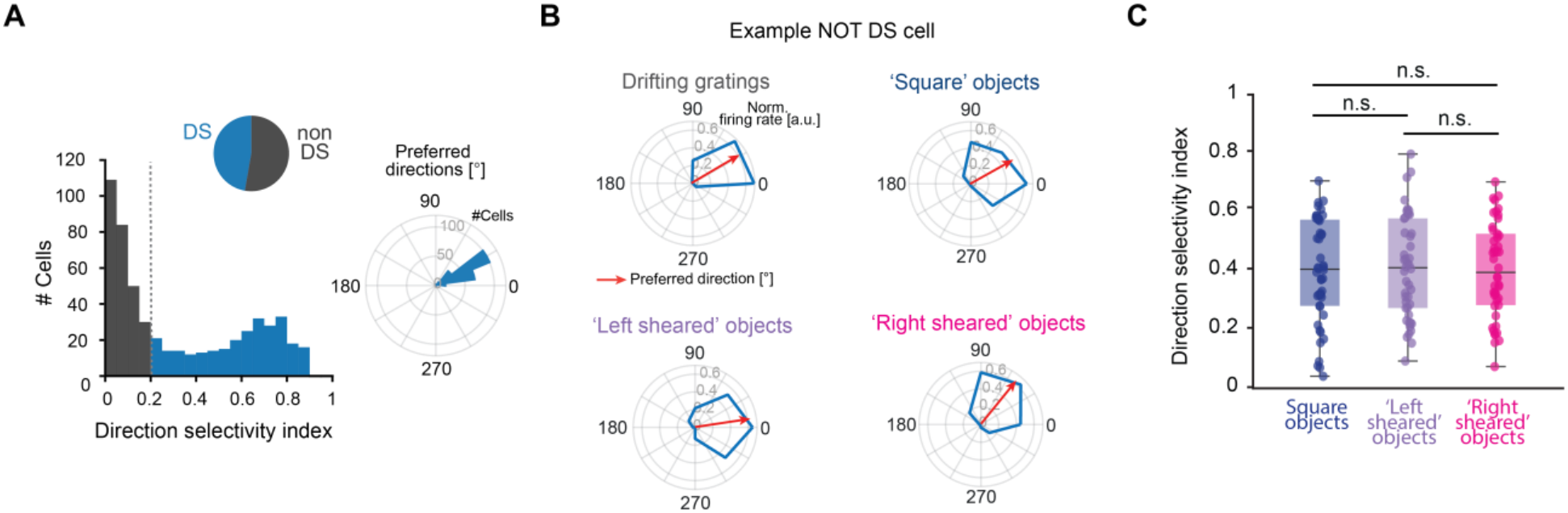
Direction selectivity of NOT cells in response to different visual stimuli. **A**, Left: distribution of direction selectivity index values in n=548 NOT cells in response to drifting gratings. Cells are deemed DS when their direction selectivity index is ≥ 0.2 (dashed line). Percentages of DS (blue) and non-direction-selective cells (non-DS, black) are shown as a pie chart in inset. Right: histogram of preferred direction of n=259 DS cells (median 25.9°). **B**, Normalized direction tuning function of one example DS NOT cell in response to drifting gratings, square and ‘left’ and ‘right sheared’ objects. Insets show one frame of the stimulus movie. Red arrow: preferred directions. **C**, Direction selectivity index distributions of n=46 cells for all object conditions tested (medians: 0.4, 0.4, 0.38). Significance shown (paired-permutation test with Bonferroni correction; P=1.000 for all comparisons).

**Fig. S3.**
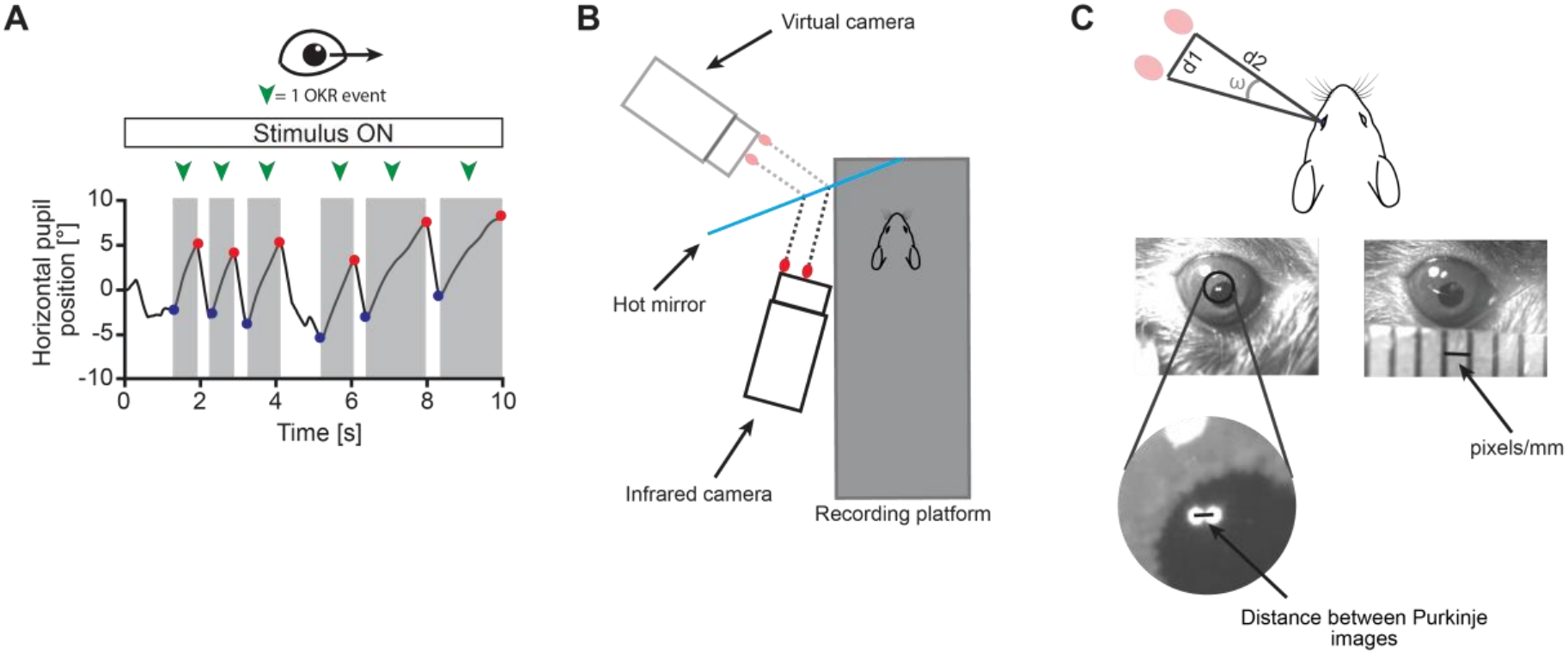
Recording of eye movements during OKR. **A**, Example of horizontal OKR for n=1 mouse and n=1 trial in response to a rightwards moving grating (0.025 cpd, 45°/s). OKR events are highlighted in gray and marked by green arrows. Red and blue dots show peaks and troughs in the pupil position over time, used to calculate the OKR gain. **B**, Schematics of recording setup. The infrared (IR) camera is placed behind the mouse (the virtual camera is its imaginary projection in the mouse frontal plane) and records eye movements from a hot mirror placed in front of the mouse. Red dots: IR LEDs. **C**, Top panel: schematics of eye position calibration: the two IR LEDs are mounted over the eye camera at a distance (d1) that can be used to calculate their subtended distance based on their virtual distance from the mouse eye (d2). Central panel: the reflected light of the LEDs (Purkinje images) is recorded from the eye video and their distance between them measured. Bottom panel: the distance between Purkinje images and the pixels/mm (x) value is used to calculate the Hirschberg ratio.

**Fig. S4.**
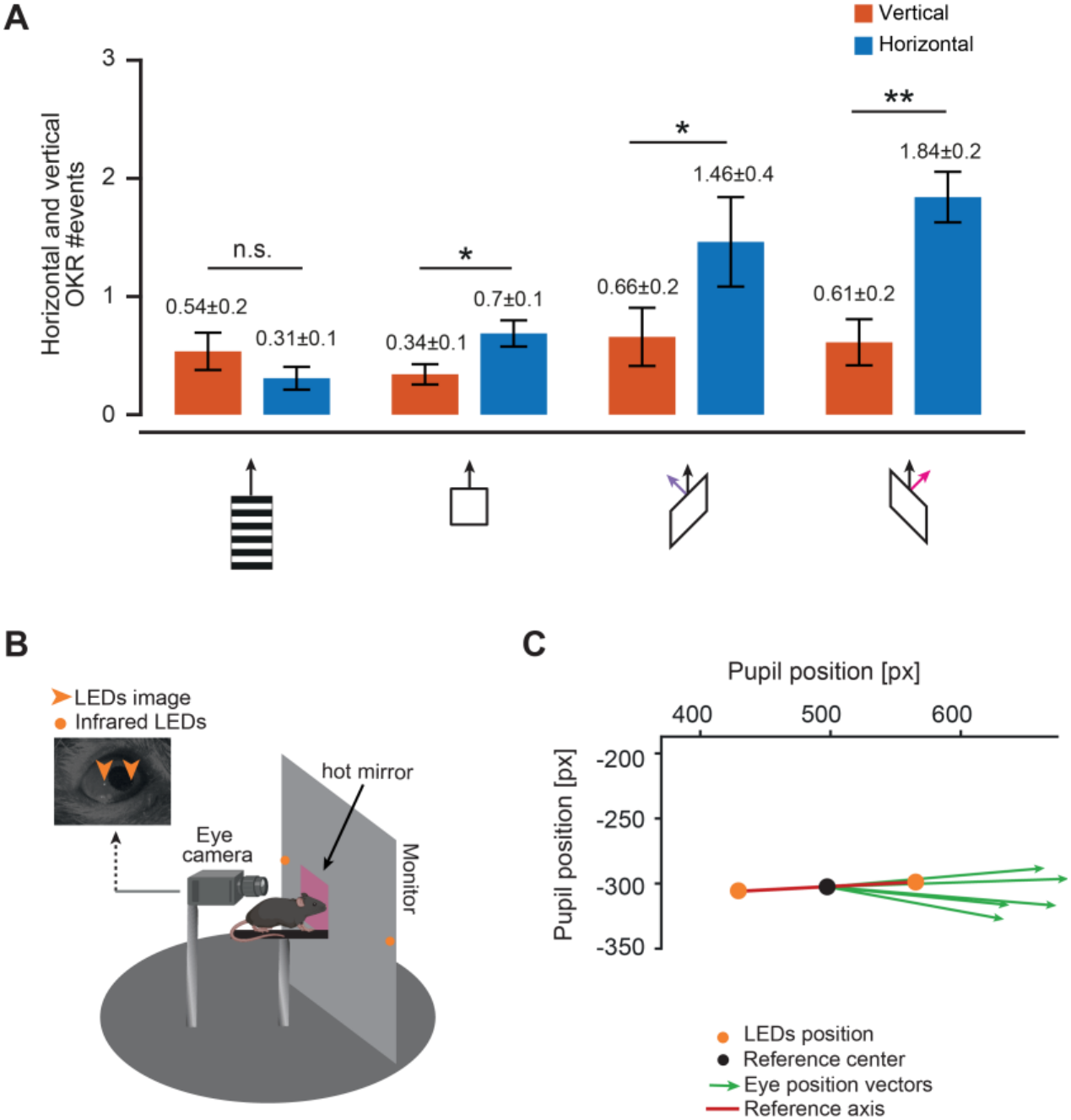
Horizontal and vertical OKR responses to vertical motion and analysis of eye movement direction. **A**, Number (mean/trial) of vertical and horizontal OKR events in response to upward-moving stimuli. Error bars: SEM over number of mice (n=8). Numbers above bars: mean±sem. Statistical difference is shown (Wilcoxon signed-rank test, left to right: P=0.062, P=0.023, P=0.0312, P=0.008). **B**, Schematics of external horizontal axis measurement, used as reference for eye movements directions. Two LEDs are mounted on the monitor and their distance measured from the eye video, marking an external reference for the horizontal meridian. **C**, Eye position vectors for each OKR event during one stimulus presentation (green arrows) plotted against the external reference axis (blue line) and with origin the center of the reference axis (black dot). Eye movement directions are calculated from the angle between eye position vectors and the reference axis.

**Fig. S5.**
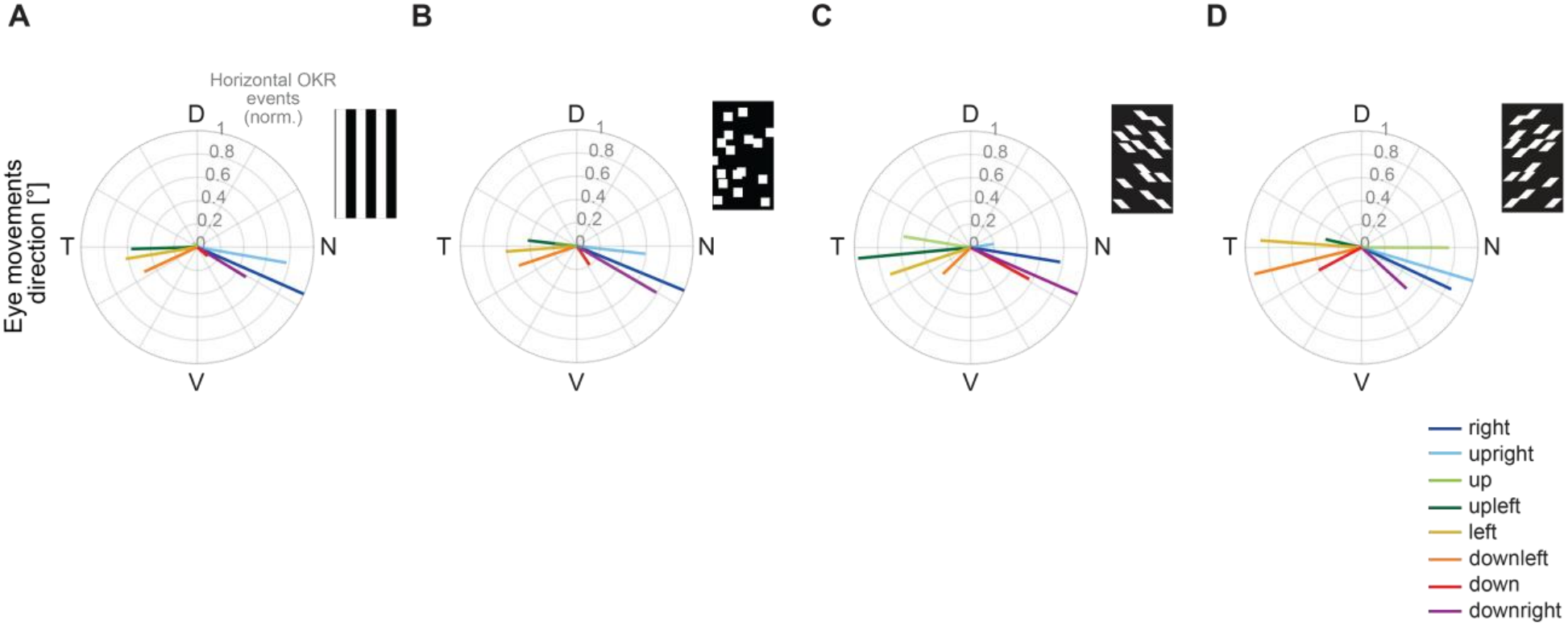
Eye movement directions across stimulus class and direction. **A-D**, Eye movement directions as a function of object direction for all stimulus conditions, color coded by motion direction, plotted as polar vectors. Vector angle indicates the subject-wise (n=6) circular mean direction, and vector lengths are normalized by the number of OKR events and normalized to global maximum.

**Fig. S6.**
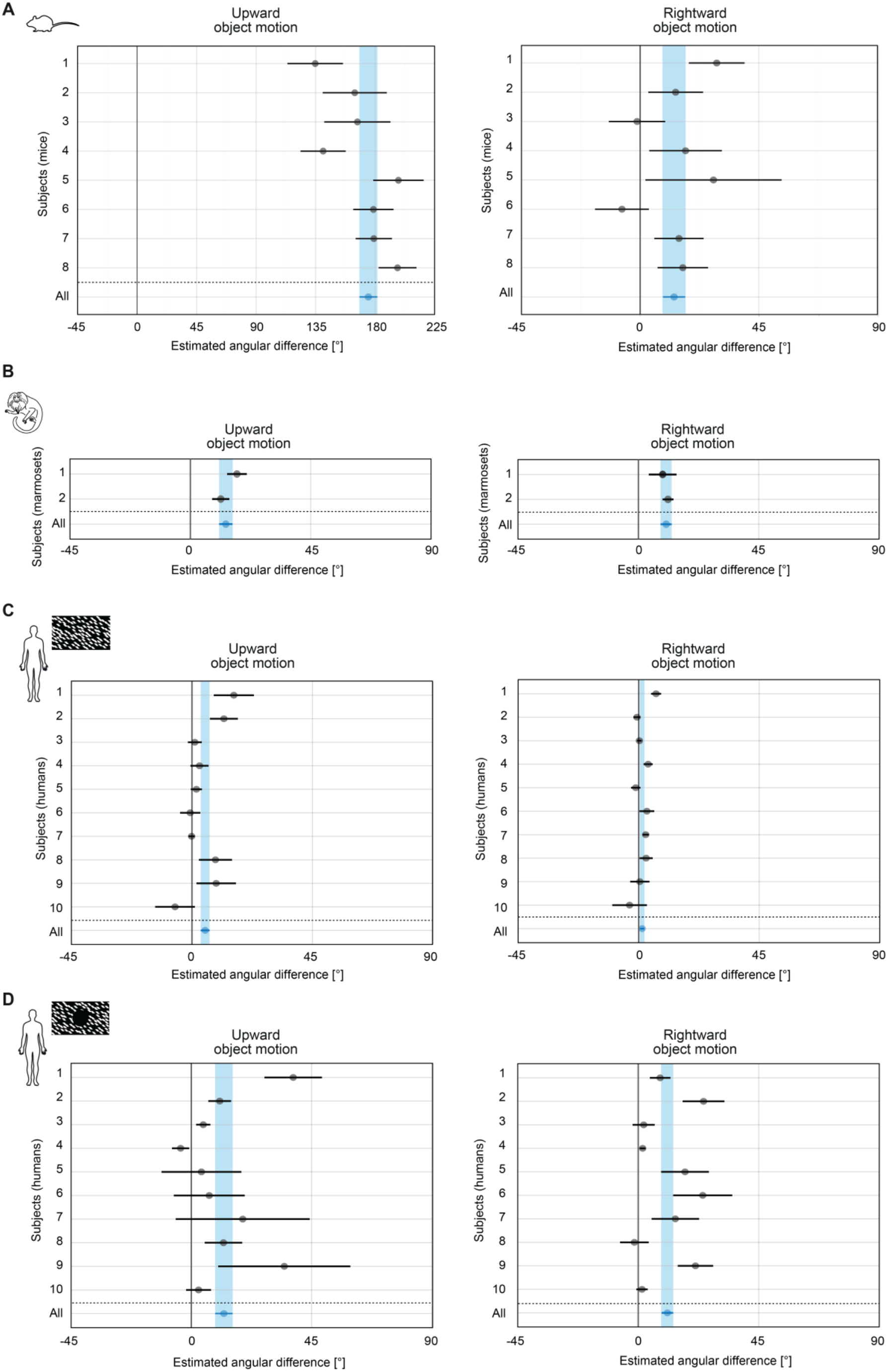
Subject-wise analysis of eye movement directions in response to upward- and rightward-moving sheared objects. **A-D**, Subject-wise confidence-interval analysis of the effect of shear condition on eye movement direction in mice (**A**, n=8), marmoset monkeys (**B**, n=2) and humans (n=10), the latter tested without (**C**) and with (**D**) a central gaze mask, for upward (left panels) and rightward (right panels) motion. Note that panel **A**, left, has a different scale on the x axis. Dots indicate subject-level effect estimates from the linear mixed-effects model, and horizontal whiskers indicate the corresponding 95% confidence intervals. The blue dots (‘All’) indicates the population-level estimates, and the blue shaded region indicates its 95% confidence interval. For upward motion in mice, signed angular estimates were wrapped to their equivalent values in the 0-360° range for visualization.

**Fig. S7.**
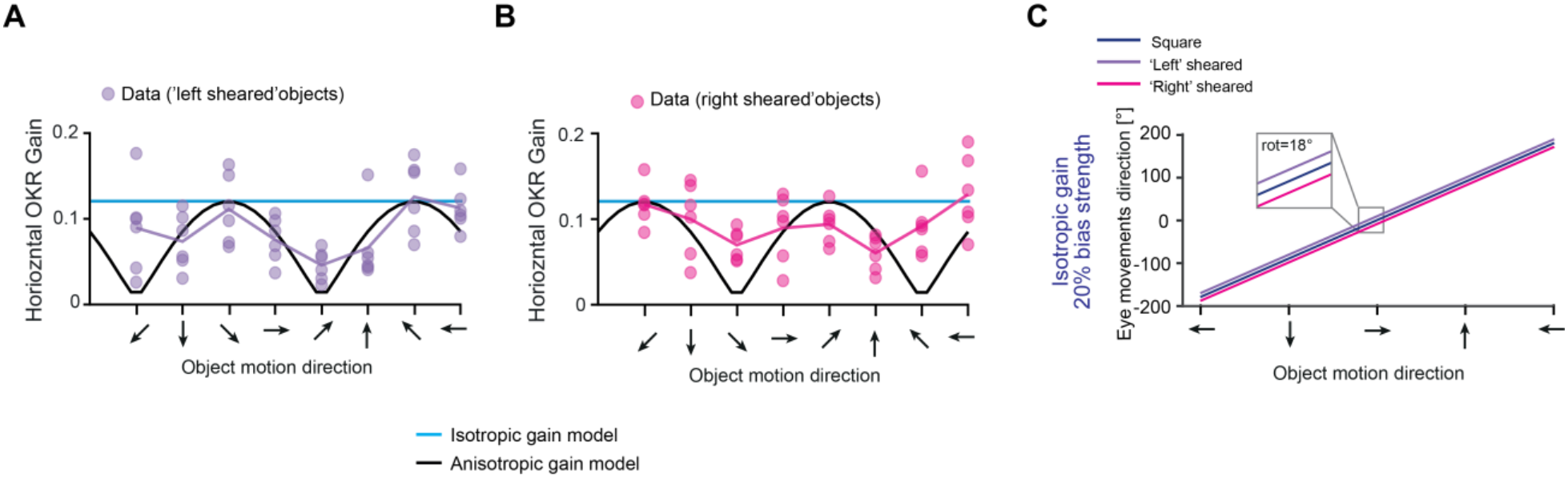
Mouse OKR responses to sheared objects predicted by isotropic and anisotropic gain models and human OKR responses predicted by an isotropic gain model. **A-B**, Gain distributions predicted by the isotropic (light blue line) and anisotropic (dark gray line) gain models as a function of global motion direction, with measured OKR gain values in response to ‘left’ and ‘right sheared’ objects (n=6 mice) overlaid. Purple and magenta continuous lines show the mean gain at each motion direction. **C**, Human eye movement direction predicted by a 20% bias strength, isotropic gain model, showing a constant shear-induced shift of 18° at each stimulus direction.

**Fig. S8.**
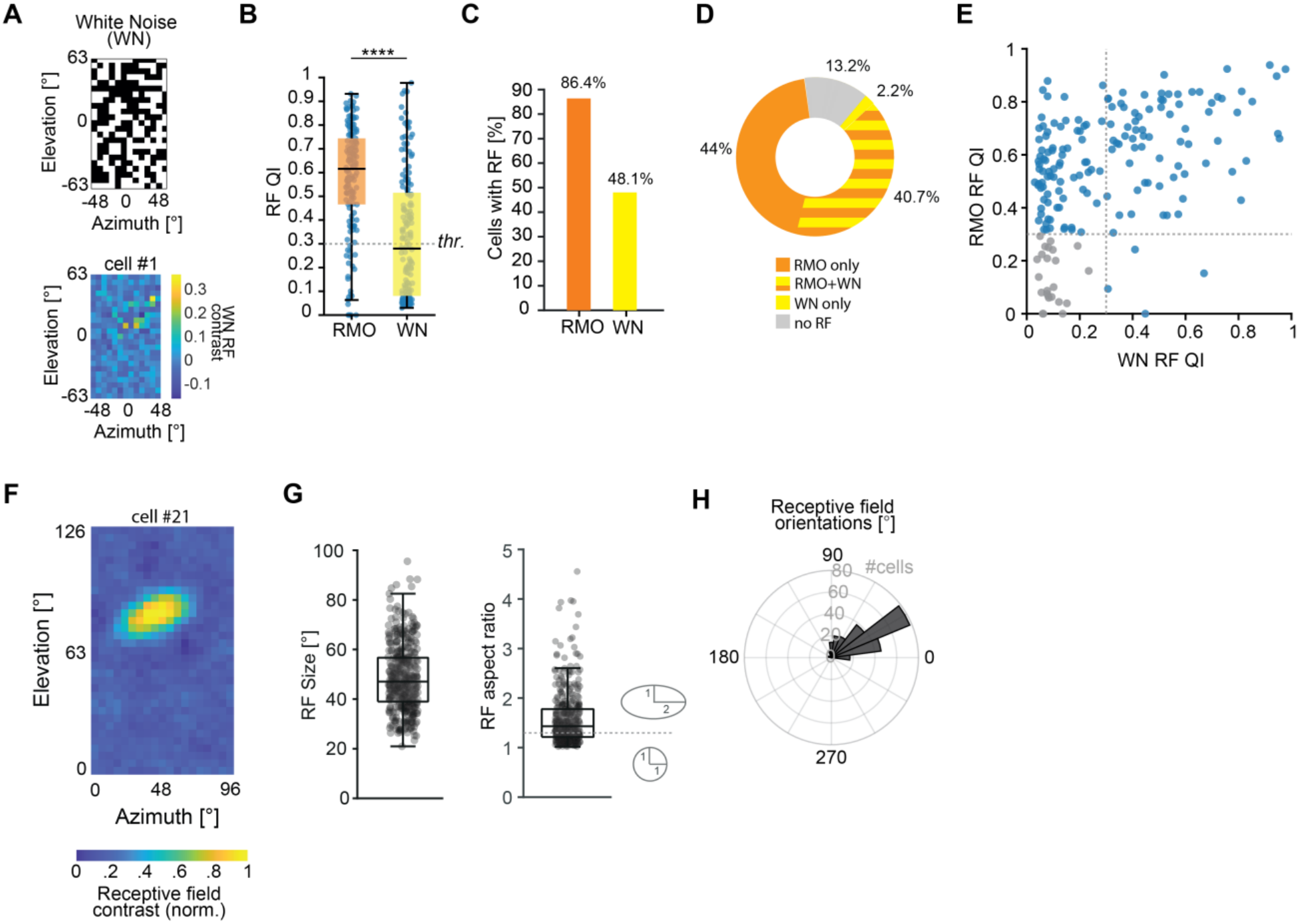
Estimation of cells’ receptive fields with white noise (WN) and random moving objects (RMOs). **A**, Top: depiction of a stimulus frame for WN on visual screen coordinates. Bottom: receptive field estimation for one exemplary DS NOT cell. **B**, Distribution of receptive fields’ quality indexes (Qi), computed based on the explained variance of a 2D gaussian fit, for n=154 NOT DS cells. Receptive fields are estimated with RMOs (orange) and WN (yellow); dashed line marks the threshold for a good receptive field (medians 0.61 and 0.28, respectively). Statistical difference (paired permutation test, p < 1 × 10^−4^). **C**, Percentage of receptive fields (Qi > 0.3) estimated with the two methods. Receptive field Qi obtained with RMO and WN. **D**, Percentage of DS cells that displayed a receptive field with RMO, WN, either stimulus (RMO+WN) and no quantifiable receptive field. **E**, Receptive field quality indices obtained with RMOs and WN for NOT DS cells; dashed lines mark Qi threshold. Blue data points have a quantifiable receptive field, gray data points are below the threshold for both RMO and WN. Panels B-E depict the same cells. **F**, Example of a receptive field for a non-DS NOT cell, estimated with the RMO stimulus and plotted as a function of visual monitor coordinates; color codes the normalized receptive field contrast. **G**, Receptive field size (left; median 47°) and aspect ratio (right; median 1.43) distributions for n=394 non-DS NOT cells. **H**, Receptive field orientation of same cells as in G (median 33.8°).

**Fig. S9.**
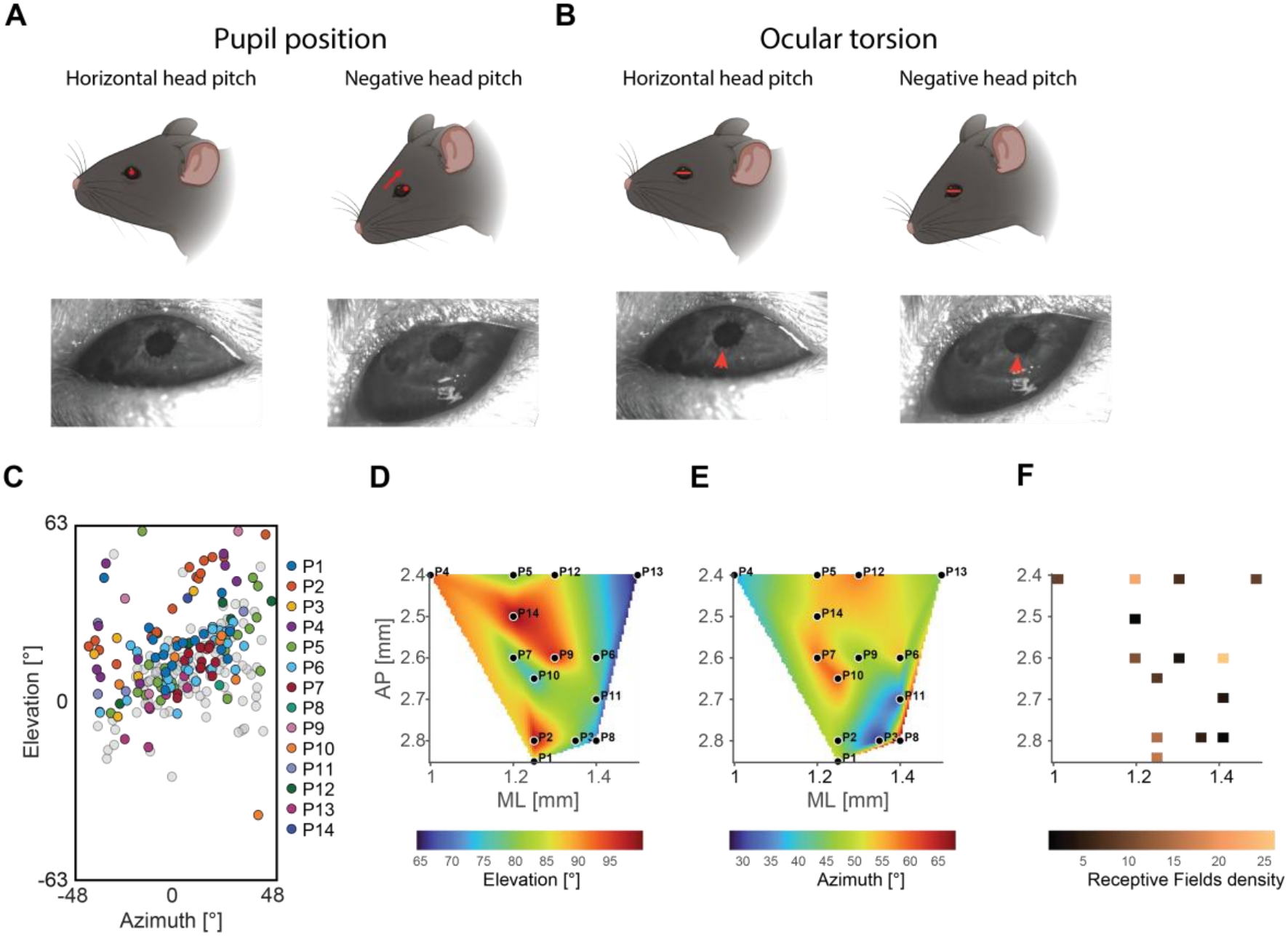
NOT receptive field organization is not explained by head position or uneven anatomical sampling. **A**, Top: Schematics of pupil position of a mouse during passive rotation of the head in the vertical (pitch) axis. Red dot is the pupil center, arrow marks the expected pupil displacement after head rotation. Bottom: Pictures of the left eye of an anesthetized mouse, showing pupil position displacement after head rotation. **B**, Top: Same as a, but illustrating ocular torsion; the red segment marks the horizontal axis of the eye. Bottom: same as a, with arrowhead overlaid marking a reference feature on the pupil used to track torsional rotation, showing that it remains stable after head rotation. **C**, Receptive field positions plotted as a function of monitor coordinates for n=137 DS cells, color-coded by recording session (electrode insertion) number (P1-P14). Gray points are cells belonging to sessions whose histology could not be validated, hence excluded from this analysis. **D**, Retinotopic map of receptive field elevation, obtained by projecting receptive field centers onto a common mediolateral-anteroposterior (ML-AP) coordinate system and interpolating elevation values using a natural-neighbor scattered interpolant (mask radius: 0.2 mm). **E**, Same as in **D**, for the azimuth axis. **F**, Receptive field density map (number of receptive fields at each recording coordinate) computed as a 2D histogram (20×20 bins) of receptive field positions in ML-AP space. Note: local extrema in the interpolated maps in **D-E** coincide with regions of lower receptive field sampling density in **F**, hence do not reflect systematic shifts in receptive field organization.

**Fig. S10.**
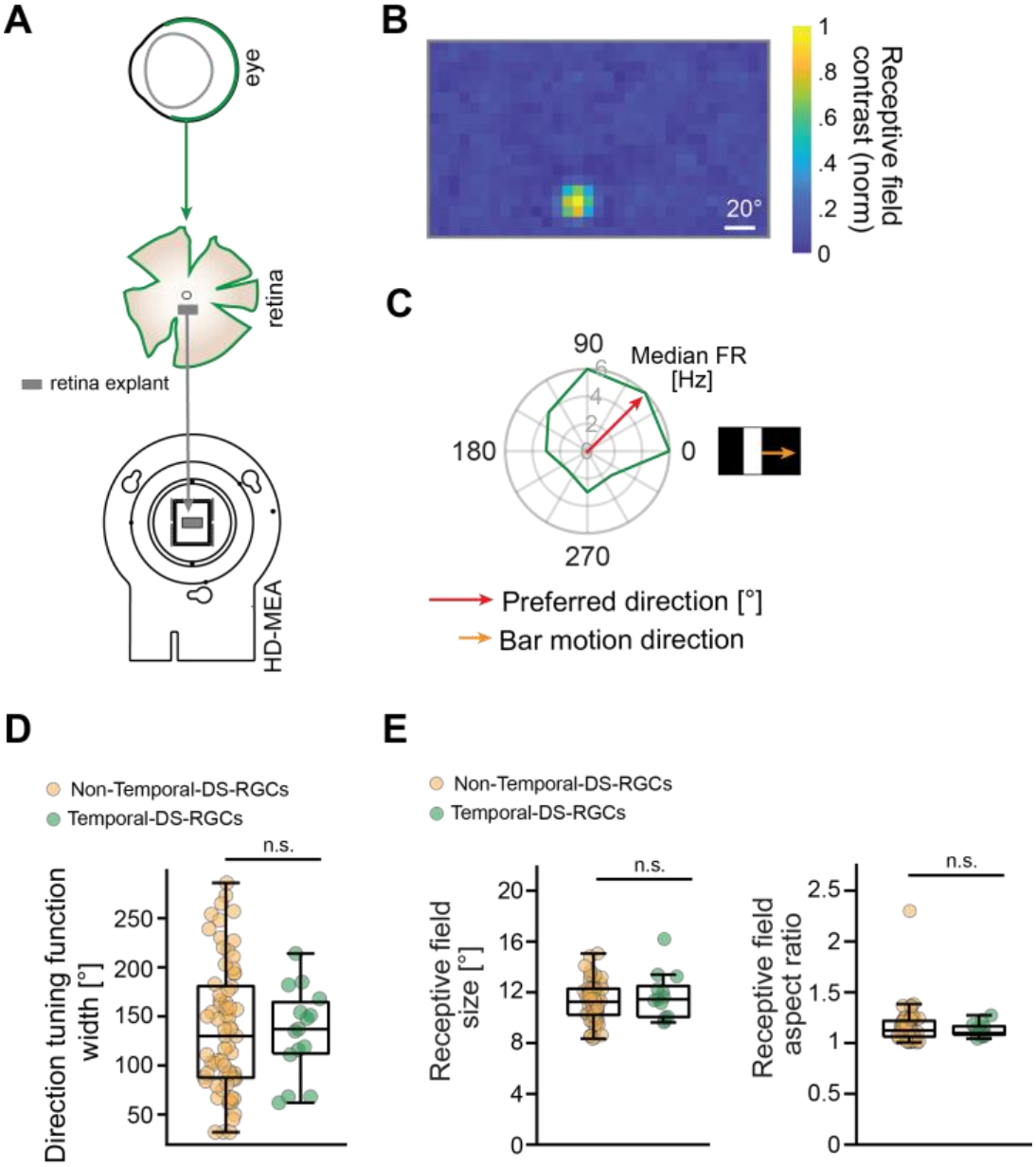
Functional characterization of DS-RGCs via HD-MEA recordings. **A**, Schematics of the ex vivo retinal preparation and recording via HD-MEA. **B**, Receptive field of an exemplary Temporal-DS-RGCs shown in HD-MEA chip coordinates (3.85 x 2.10 mm^2^). Color codes the normalized receptive field contrast. **C**, Direction tuning function, in visual coordinates, and preferred direction for a Temporal-DS-RGCs in response to a moving bar (depicted in inset, shown at a size of 20° and speed of 10 °/s). **D**, Distribution of direction tuning curve widths, quantified as the circular full width at half maximum (FWHM) of each cell’s direction tuning curve, for non-Temporal DS-RGCs (orange, n=69) and Temporal-DS-RGCs (green, n=15). Median (dashed lines): 130° and 137°, respectively. Statistical difference is reported (Ranksum test, P=0.921). **E**, Distributions of receptive field sizes (left panel) and aspect ratios (right panel) of non-Temporal DS-RGCs (orange, n=64) and Temporal-DS-RGCs (green, n=13) with Qi ≥ 0.3. Median size, 11.3° and 11.5°, respectively. Median aspect ratio 1.12 and 1.1, respectively. Statistical difference is reported (Ranksum test, P=0.62 and P=0.739, respectively).

**Fig. S11.**
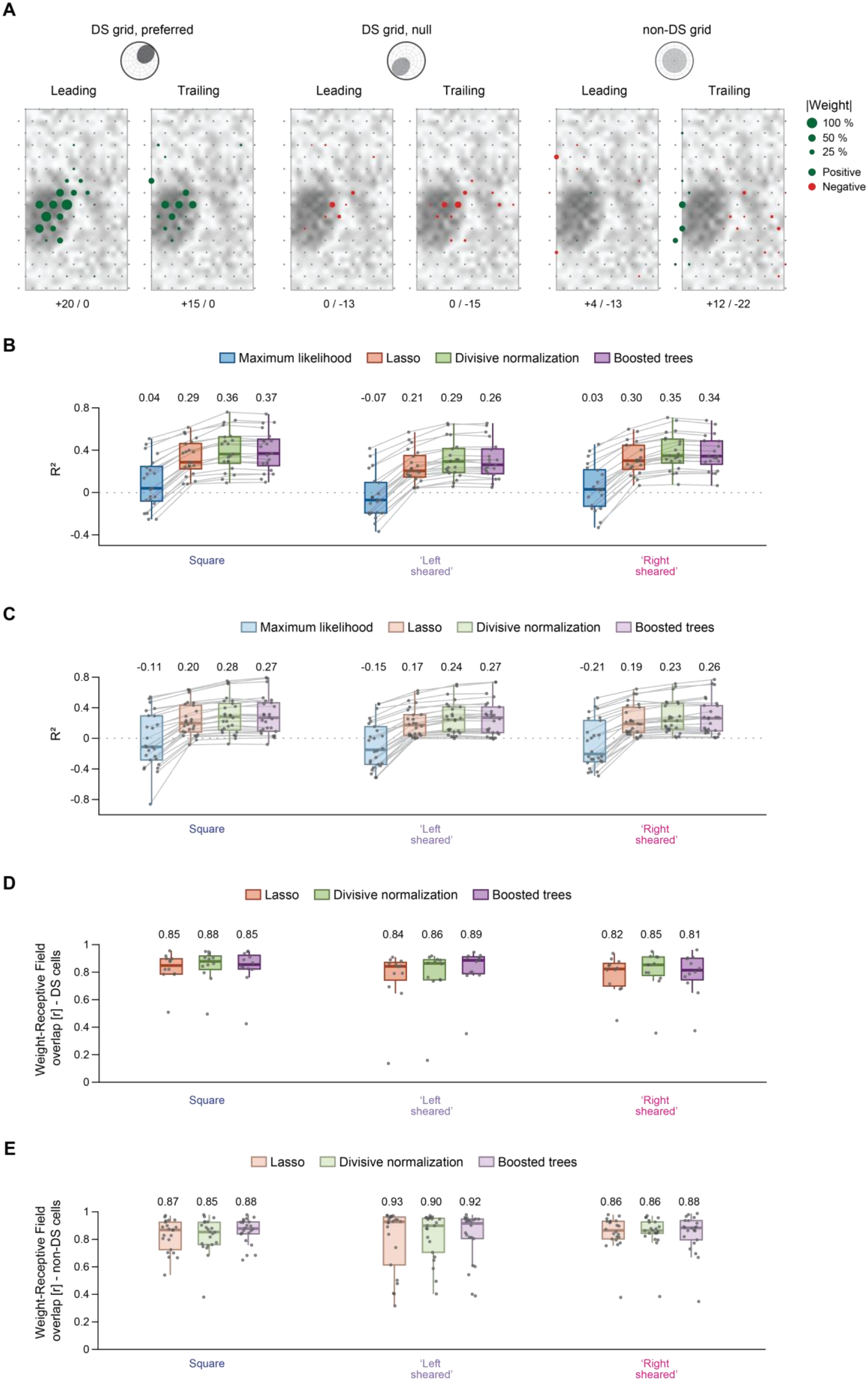
Spatial maps of model weight for the leading- and trailing-edge, model accuracy, and alignment of weights with receptive fields across models. **A**, Weights of the example cell of Fig. 5B (square objects), each grid shown separately for the leading and the trailing edge. Dot area is proportional to the weight magnitude, scaled to the largest single-edge node weight of the six maps (100%); green, positive; red, negative; background as in Fig. 5B. Numbers below the maps: number of grid nodes (of 136) with a positive/negative weight. **B**, Cross-validated R^2^ of DS NOT cells (n = 21) for four NOT-stage models. All four share the fixed retinal stage and the fixed output rectification of Fig. 5A: a linear model fitted without regularization (Maximum likelihood); a linear model fitted with L1 regularization (Lasso); the same L1-regularized model fitted to divisively normalized inputs (Divisive normalization, the model shown in Fig. 5); and boosted regression trees. Only the divisive normalization model used normalized inputs. Without regularization, the linear model did not generalize to held-out data (median R^2^ −0.07 to 0.04), indicating that the fitted weights might have been underdetermined by the data. All other models were therefore regularized, either by an L1 penalty or, for boosted trees, by the cross-validated number of trees. Lines connect the values of each cell; medians are annotated above boxes. Boxes as in Fig. 5F. **C**, Same as B for non-DS NOT cells (n = 26). **D**, Alignment of model weights and receptive fields for Lasso, Divisive normalization, Boosted trees models in DS NOT cells (n = 12). Pearson correlation between each cell’s weight-magnitude map and the Gaussian fit of its measured receptive field for square, ‘right sheared’ and ‘left sheared’ objects. For boosted trees each input’s feature importance was used instead. Only cells with a receptive field quality index Qi > 0.3 are included. Median values are annotated above boxes. **E**, Same as D for non-DS NOT cells (n = 22).

**Table S1.**
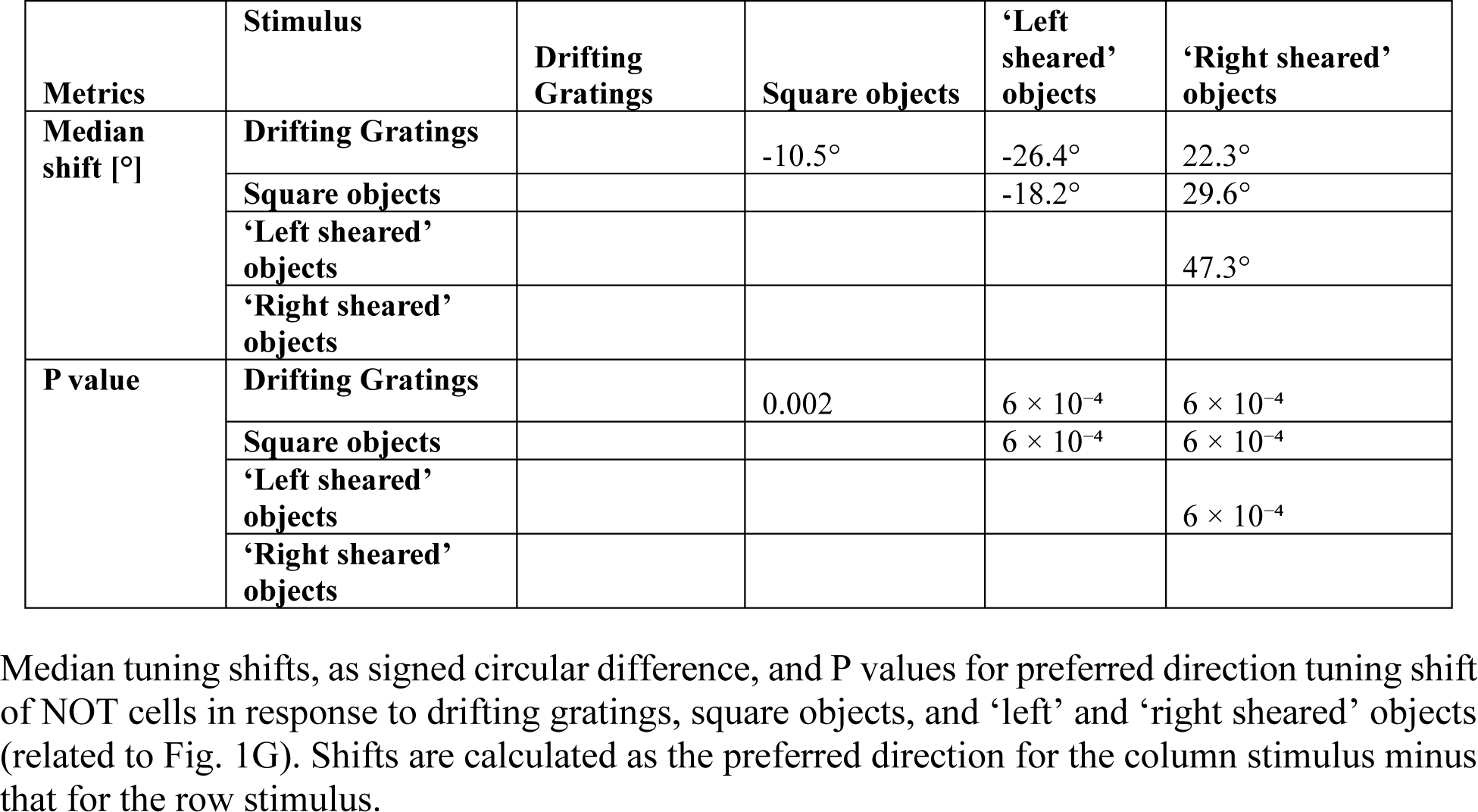

| **Metrics** | **Stimulus** | **Drifting Gratings** | **Square objects** | **‘Left sheared’ objects** | **‘Right sheared’ objects** |
|---|---|---|---|---|---|
| **Median shift [°]** | **Drifting Gratings** |  | -10.5° | -26.4° | 22.3° |
|  | **Square objects** |  |  | -18.2° | 29.6° |
|  | **‘Left sheared’ objects** |  |  |  | 47.3° |
|  | **‘Right sheared’ objects** |  |  |  |  |
| **P value** | **Drifting Gratings** |  | 0.002 | 6 × 10<sup>−4</sup> | 6 × 10<sup>−4</sup> |
|  | **Square objects** |  |  | 6 × 10<sup>−4</sup> | 6 × 10<sup>−4</sup> |
|  | **‘Left sheared’ objects** |  |  |  | 6 × 10<sup>−4</sup> |
|  | **‘Right sheared’ objects** |  |  |  |  |

**Table S2.**

| **Motion direction** | **Subject number** | ***rot* (°)** | **β (°)** | **95% CI** | **P value** |
|---|---|---|---|---|---|
| **Rightward** | 01 | 29.3 | 29.0 | 18.5, 39.6 | 1.90 × 10<sup>−6</sup> |
|  | 02 | 13.3 | 13.5 | 3.08, 23.9 | 0.012 |
|  | 03 | -1.2 | -1.17 | -11.8, 9.52 | 0.827 |
|  | 04 | 15.3 | 17.2 | 3.48, 31.0 | 0.015 |
|  | 05 | 26.1 | 27.8 | 1.98, 53.6 | 0.036 |
|  | 06 | -6.7 | -6.9 | -17.1, 3.31 | 0.179 |
|  | 07 | 14.7 | 14.7 | 5.38, 24.1 | 0.003 |
|  | 08 | 16.2 | 16.2 | 6.63, 25.7 | 0.001 |
|  | Population | 13.4 | 12.9 | 8.64, 17.1 | 5.61 × 10<sup>−9</sup> |
| **Upward** | 01 | 133.4 | 135 | 114, 156 | 4.12 × 10<sup>−16</sup> |
|  | 02 | 165.3 | 165 | 140, 189 | 4.13 × 10<sup>−12</sup> |
|  | 03 | 158.6 | 167 | 142, 192 | 2.38 × 10<sup>−16</sup> |
|  | 04 | 140.7 | 141 | 124, 158 | 1.39 × 10<sup>−13</sup> |
|  | 05 | 198.0 | 198 | 179, 217 | 6.82 × 10<sup>−17</sup> |
|  | 06 | 178.5 | 179 | 164, 194 | 7.35 × 10<sup>−21</sup> |
|  | 07 | 179.0 | 179 | 165, 193 | 6.80 × 10<sup>−25</sup> |
|  | 08 | 195.4 | 197 | 183, 211 | 5.07 × 10<sup>−25</sup> |
|  | Population | 168.6 | 175 | 168, 182 | 1.19 × 10<sup>−132</sup> |

**Table S3.**

| **Motion direction** | **Subject number** | ***rot* (°)** | **β (°)** | **95% CI** | **P value** |
|---|---|---|---|---|---|
| **Rightward** | 01 | 9.2 | 9.07 | 3.90–14.2 | 6.41 × 10<sup>−4</sup> |
|  | 02 | 11.3 | 11.1 | 9.05–13.1 | 1.80 × 10<sup>−24</sup> |
|  | Population | 10.2 | 10.4 | 8.29–12.5 | 3.17 × 10<sup>−21</sup> |
| **Upward** | 01 | 17.3 | 17.4 | 13.7–21.1 | 2.53 × 10<sup>−17</sup> |
|  | 02 | 12 | 11.4 | 8.14–14.6 | 1.84 × 10<sup>−11</sup> |
|  | Population | 14.6 | 13.3 | 10.8–15.8 | 1.21 × 10<sup>−23</sup> |

**Table S4.**

| **Motion direction** | **Subject number** | ***rot* (°)** | **β (°)** | **95% CI** | **P value** |
|---|---|---|---|---|---|
| **Rightward** | 01 | 6.5 | 6.49 | 4.63, 8.36 | 2.84 × 10<sup>−8</sup> |
|  | 02 | -0.6 | -0.644 | -1.95, 0.665 | 0.332 |
|  | 03 | 0.3 | 0.259 | -0.580, 1.10 | 0.527 |
|  | 04 | 3.5 | 3.52 | 1.83, 5.21 | 1.03 × 10<sup>−4</sup> |
|  | 05 | -1.1 | -1.10 | -2.86, 0.650 | 0.215 |
|  | 06 | 3.2 | 3.05 | 0.305, 5.80 | 0.0297 |
|  | 07 | 2.6 | 2.62 | 1.43, 3.81 | 3.22 × 10<sup>−5</sup> |
|  | 08 | 2.8 | 2.78 | 0.327, 5.23 | 0.027 |
|  | 09 | 0.8 | 0.424 | -3.18, 4.03 | 0.816 |
|  | 10 | -3.3 | -3.40 | -9.86, 3.07 | 0.293 |
|  | Population | 2.5 | 1.31 | 0.467, 2.15 | 0.002 |
| **Upward** | 01 | 15.8 | 15.6 | 8.16, 23.1 | 9.54 × 10<sup>−5</sup> |
|  | 02 | 11.8 | 11.9 | 6.74, 17.1 | 1.56 × 10<sup>−5</sup> |
|  | 03 | 1.1 | 1.08 | -1.51, 3.66 | 0.402 |
|  | 04 | 2.8 | 2.83 | -0.561, 6.21 | 0.100 |
|  | 05 | 1.7 | 1.66 | -0.416, 3.73 | 0.116 |
|  | 06 | -0.6 | -0.643 | -4.41, 3.12 | 0.736 |
|  | 07 | -0.1 | -0.120 | -1.26, 1.02 | 0.834 |
|  | 08 | 8.3 | 8.78 | 2.59, 15.0 | 0.006 |
|  | 09 | 8.7 | 9.06 | 1.70, 16.4 | 0.017 |
|  | 10 | -6.1 | -6.31 | -13.7, 1.12 | 0.093 |
|  | Population | 5.7 | 4.94 | 3.28, 6.59 | 6.85 × 10<sup>−9</sup> |

**Table S5.**
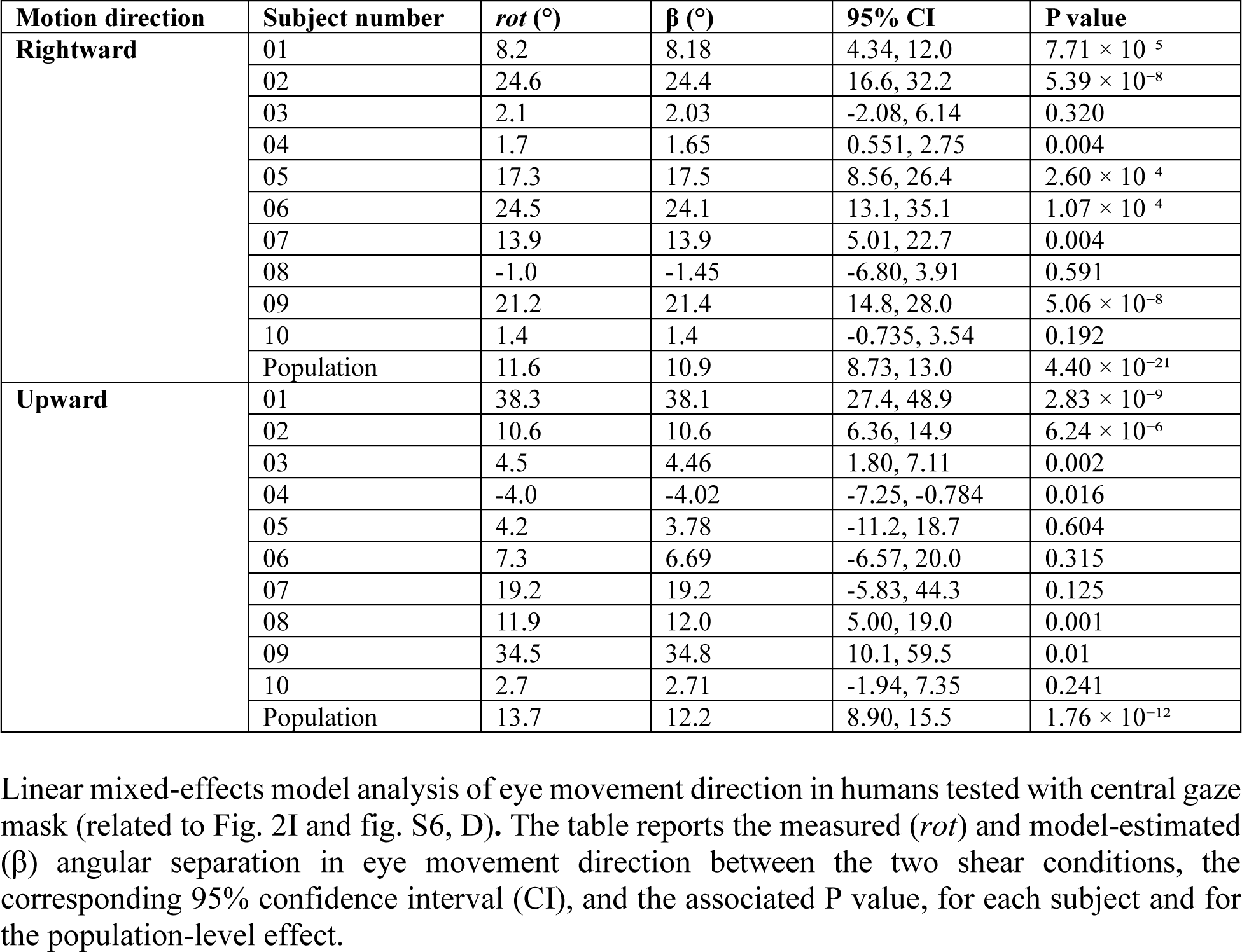

**Table S6.**

| **Motion direction** | **Subject number** | **β (°)** | **95% CI** | **P value** |
|---|---|---|---|---|
| Rightward | 01 | 1.68 | −3.19, 6.56 | 0.494 |
|  | 02 | 25.0 | 19.6, 30.5 | 1.76 × 10<sup>−16</sup> |
|  | 03 | 1.77 | −3.03, 6.57 | 0.461 |
|  | 04 | −1.86 | −3.89, 0.159 | 0.0706 |
|  | 05 | 18.6 | 12.6, 24.5 | 3.84 × 10<sup>−9</sup> |
|  | 06 | 21.0 | 13.3, 28.7 | 2.29 × 10<sup>−7</sup> |
|  | 07 | 11.2 | 6.71, 15.8 | 3.36 × 10<sup>−6</sup> |
|  | 08 | −4.22 | −9.30, 0.849 | 0.102 |
|  | 09 | 20.9 | 14.1, 27.8 | 1.31 × 10<sup>−8</sup> |
|  | 10 | 4.80 | −1.61, 11.2 | 0.140 |
|  | **Population** | **9.80** | **7.84, 11.8** | **6.08 × 10<sup>−22</sup>** |
| Upward | 01 | 22.5 | 9.74, 35.3 | 6.82 × 10<sup>−4</sup> |
|  | 02 | −1.32 | −8.74, 6.10 | 0.726 |
|  | 03 | 3.38 | −0.284, 7.04 | 0.0699 |
|  | 04 | −6.84 | −11.5, −2.21 | 0.00412 |
|  | 05 | 2.12 | −5.71, 9.95 | 0.593 |
|  | 06 | 7.34 | −2.68, 17.4 | 0.150 |
|  | 07 | 19.4 | 8.24, 30.5 | 7.97 × 10<sup>−4</sup> |
|  | 08 | 3.23 | −6.94, 13.4 | 0.532 |
|  | 09 | 25.7 | 6.72, 44.7 | 0.00865 |
|  | 10 | 9.02 | −0.240, 18.3 | 0.0560 |
|  | **Population** | **7.49** | **4.17, 10.8** | **1.04 × 10<sup>−5</sup>** |

**Table S7.**

| **Metric** | **Stimulus position** | **1** | **2** | **3** | **4** | **FF** |
|---|---|---|---|---|---|---|
| **Median FR** | **1** |  | 0.052 | 0.005 | 0.006 | 0.002 |
|  | **2** | 0.052 |  | 0.004 | 0.133 | 0.007 |
|  | **3** | 0.005 | 0.004 |  | 0.002 | 1.000 |
|  | **4** | 0.006 | 0.133 | 0.002 |  | 0.004 |
|  | **FF** | 0.002 | 0.007 | 1.000 | 0.004 |  |
| **Gain** | **1** |  | 0.937 | 0.045 | 0.045 | 0.045 |
|  | **2** | 0.937 |  | 0.045 | 0.078 | 0.045 |
|  | **3** | 0.045 | 0.045 |  | 0.045 | 1.000 |
|  | **4** | 0.045 | 0.078 | 0.045 |  | 0.045 |
|  | **FF** | 0.045 | 0.045 | 1.000 | 0.045 |  |
| **Number of events** | **1** |  | 0.531 | 0.045 | 0.045 | 0.045 |
|  | **2** | 0.531 |  | 0.045 | 0.117 | 0.045 |
|  | **3** | 0.045 | 0.045 |  | 0.045 | 0.521 |
|  | **4** | 0.045 | 0.117 | 0.045 |  | 0.045 |
|  | **FF** | 0.045 | 0.045 | 0.521 | 0.045 |  |

**Table S8.**

| **Subject number** | **Gender** | **Age** | **Visual acuity by autorefraction** |
|---|---|---|---|
| 01 | M | 43 | OD:1.25 OS:1.25 |
| 02 | F | 31 | OD:1.0 OS:1.0 |
| 03 | M | 35 | OD:1.1 OS:1.1 |
| 04 | F | 32 | OD:0.8 OS:1.0 |
| 05 | M | 38 | OD:1.0 OS:1.0 |
| 06 | M | 37 | OD:1.0 OS:1.0 |
| 07 | M | 32 | OD:1.25 OS:1.25 |
| 08 | M | 29 | OD:1.0 OS:1.0 |
| 09 | F | 37 | OD:1.25 OS:1.25 |
| 10 | M | 41 | OD:1.0 OS:1.0 |

## Ethical statement

All animal procedures and experiments were performed in accordance with the Swiss Federal and Animal Protection Law and authorized by the Cantonal Veterinary Office (Basel, Switzerland) under the license number 3150 (authorization number 37133), 3071 (authorization number 35545) and 3236 (authorization number 36609).

All experimental procedures with marmoset monkeys followed the National Institutes of Health Guide for the Care and Use of Laboratory Animals. The protocol for experimental and behavioral procedures (#101677/2015-008) was approved by the University of Rochester Institutional Animal Care and Use Committee.

All experiments with human participants were conducted in collaboration with the Augenklinik of the Universitätsspital Basel within the IOB-EYEConic-001 research study, in accordance with Swiss legal requirements and the World Medical Association Declaration of Helsinki, and were approved by the Ethikkommission Nordwest-und Zentralschweiz (EKNZ; project ID: 2025-01825). All participants provided written informed consent prior to participation.

## Acknowledgements

We thank Catalin Mitelut, Robin Tremblay and Zoltan Raics for their support in setting up the experimental rig and the acquisition software. We are grateful to Nils Schaerer, Ursula Hall, Tiana Koottungal, Mirella Barboni, Daniela Hauenstein and Lucas Janeschitz-Kriegl, for their guidance and support in the EYEConic study, and to all the participants, whose kind contributions have greatly advanced our work and scientific knowledge. We thank Frank Schaeffel for his support with eye tracking data collection and analysis. We thank Marco Cattaneo for his support with the statistical analysis. We thank Alex Fratzl, Tamas Dalmay, Alberto Pauletti, Dominique Siegenthaler, Igor Frangez and Botond Roska for their feedback during manuscript preparation. We thank Chiara Pivetta, Karina Drumm, and the staff of the IOB animal facility for their support with animal care and experimental protocols.

## Author contributions

Conceptualization: FF, FBR, MB. Data curation: FBR, MB, AB. Formal analysis: MB, FBR, FF. Funding acquisition: FBR, JFM, FF. Investigation: FBR, MB, AB, JFM, FF. Methodology: FBR, MB, FF. Project administration: FF. Resources: FF. Software: MB, FBR, AB, JFM, FF. Supervision: FF, FBR. Validation: FBR, MB. Visualization: FBR, MB, FF. Writing - original draft: FBR, FF. Writing - review & editing: FBR, MB, JFM, FF, AB.

